# Estrogen receptor alpha regulates alcohol craving in females: convergent evidence from mouse models and human genetics

**DOI:** 10.64898/2026.06.03.729849

**Authors:** Roberto Pagano, Lali Kruashvili, Monika Puchalska, Liubov S. Kalinichenko, Yamna Rizwan, Julia Świderska, Bartosz Wojtas, Bartlomiej Gielniewski, Elena Choleris, Jerzy Samochowiec, Swapnil Awasthi, Patrick Bach, Josef Frank, Andreas Heinz, Sabine Hoffmann, Stephan Ripke, Michael Smolka, Stephanie H. Witt, Christiane Mühle, Johannes Kornhuber, Falk Kiefer, Christian P. Müller, Bernd Lenz, Kasia Radwanska

**Affiliations:** Nencki Institute of Experimental Biology of Polish Academy of Sciences, 3 Pasteur St., Warsaw 02-093, Poland; Laboratory of Molecular and Cellular Neurobiology, International Institute of Molecular and Cell Biology, Warsaw, Poland; Department of Psychiatry and Psychotherapy, Friedrich-Alexander University Erlangen-Nürnberg, Erlangen, Germany; Department of Psychology and Neuroscience Program, University of Guelph, Guelph, Ontario, Canada; Pomeranian Medical University, Szczecin, Poland; Research Division Experimental Psychiatry and (Epi)Genomics, Department of Psychiatry and Psychotherapy, Charité – University Hospital Berlin, Berlin, Germany; Department of Addictive Behavior and Addiction Medicine, Central Institute of Mental Health, Medical Faculty Mannheim, Heidelberg University, Mannheim, Germany; Department of Genetic Epidemiology in Psychiatry, Central Institute of Mental Health, Medical Faculty Mannheim, Heidelberg University, Mannheim, Germany; Department of Psychiatry and Psychotherapy, University of Tuebingen, Tuebingen, Germany; Department of Psychiatry, Technische Universität Dresden, Dresden, Germany; Biobank of the Center for Innovative Psychiatric and Psychotherapeutic Research, Central Institute of Mental Health, Medical Faculty Mannheim, Heidelberg University, Mannheim, Germany; Institute of Psychopharmacology, Central Institute of Mental Health, Medical Faculty Mannheim, Heidelberg University, Mannheim, Germany

## Abstract

Alcohol craving and consumption fluctuate across the reproductive cycle in females with alcohol use disorder (AUD), suggesting that estrogen signaling contributes to disease vulnerability. Here, we investigated the role of estrogen receptor alpha (ERα; *Esr1*; *ESR1*) in alcohol seeking using complementary mouse and human approaches, as this receptor was previously identified as a risk factor for AUD. Female mice were characterized in an IntelliCage-based multidimensional AUD paradigm that stratifies individuals into AUD-prone and AUD-resistant phenotypes. Transcriptomic profiling of the amygdala revealed that Esr1 is a top transcription factor for differentially expressed genes in mice drinking alcohol, and the estrogen signaling pathway was deregulated specifically in AUD-prone mice. Although alcohol exposure did not alter overall *Esr1*/ERα mRNA or protein abundance, both transcript and protein levels positively correlated with cue-induced alcohol seeking, indicating that inter-individual variation in ERα signaling predicts relapse-like behavior. Causal manipulations confirmed a functional role of ERα. Local knockdown of *Esr1* in the basolateral amygdala reduced excitatory synaptic transmission, attenuated alcohol motivation, cue-induced seeking, and relapse drinking, and impaired cue-associated memory recall without affecting anxiety-like behavior. Similarly, ovariectomy decreased amygdala ERα expression, altered synaptic protein markers, and reduced alcohol-seeking behaviors, supporting regulation by endogenous ovarian hormones. Extending these findings to humans, *ESR1* gene polymorphisms (rs6902771, rs11155819 and rs6557171) were associated with the probability of alcohol binge drinking and alcohol consumption days as well as craving and loss of control in real world in a longitudinal clinical cohort, while *ESR1* mRNA blood levels were increased in women with AUD diagnosis. Together, these convergent molecular, circuit, behavioral, and genetic data identify ERα signaling in the amygdala as an important modulator of alcohol-seeking behavior induced by alcohol cue and relapse vulnerability, highlighting estrogen pathways as potential therapeutic targets and markers for AUD.

## INTRODUCTION

Alcohol use disorder (AUD) is a chronic, progressive psychiatric condition with limited therapeutic options that benefit only a subset of patients [1–3]. This unmet clinical need calls for a deeper mechanistic understanding of the disease and the development of more effective, personalized treatments. Historically, addiction research has focused predominantly on males, reflecting the higher prevalence of substance abuse in men and resulting in a substantial gap in our understanding of female-specific mechanisms and treatments for AUD [4]. However, recent data indicate that the gap in binge drinking frequency between men and women has narrowed considerably [5]. Moreover, accumulating clinical and preclinical evidence demonstrates that females exhibit the so-called *telescoping effect*, characterized by a more rapid transition from initial alcohol use to dependence, more severe withdrawal symptoms, and higher relapse rates [6–8]. Together, these findings suggest the existence of female-specific biological mechanisms underlying AUD vulnerability.

Consistent with this view, fluctuations in estrogens bioavailability across the menstrual cycle predict alcohol craving and consumption in women with AUD [9, 10]. The likelihood of binge drinking is significantly higher during the ovulatory phase, when 17β-estradiol (E2) levels peak and the progesterone-to-estradiol ratio is low [9, 10]. Animal studies further support a role for ovarian hormones in modulating alcohol consumption. Female rodents consume more alcohol during estrus and proestrus, phases associated with elevated E2 levels, compared with diestrus and metestrus [11–13]. In addition, alcohol rewarding effects are diminished in ovariectomized animals and reduced alcohol intake following ovariectomy can be restored by estrogen replacement [13, 14]. Collectively, these findings implicate estrogen signaling as an important modulator of alcohol-related behaviors in females.

The behavioral effects of estrogen are mediated primarily by estrogen receptors α (ERα) and β (ERβ) as well as G Protein-Coupled ER (GPER), which are widely expressed in brain regions implicated in addiction [15–19]. In the present study, we focused on ERα because of its established roles in regulating excitatory neurotransmission [20], promoting long-term potentiation (LTP) [21, 22], and coupling with metabotropic glutamate receptors (mGluR1/5) [23], thereby activating diverse signaling cascades that influence synaptic plasticity and animal behavior [24, 25]. In the context of AUD, ERα activation was demonstrated to enhance alcohol-induced firing of dopaminergic neurons in the ventral tegmental area (VTA) and increase alcohol consumption more than activation of ERβ [26]. Also blocking ERα, but not ERβ, in the bed nucleus of the stria terminalis (BNST) abolished the pro-alcohol drinking effects of high ovarian E2 status [27], while brain-penetrant selective ERα degrader (SERD), YL3-122, decreased binge-like alcohol drinking [28]. Moreover, genetic studies have linked several single nucleotide polymorphisms (SNPs) in the *ESR1* gene, which encodes ERα, to alcohol dependence [9, 29]. Despite these observations, the contribution of ERα-dependent brain mechanisms to AUD-related behaviors beyond alcohol intake remains unexplored.

To address this gap, we investigated the role of ERα in the amygdala in regulating AUD-related behaviors. We focused on the amygdala for two main reasons. First, ERα is highly expressed in this region [16, 19, 30]. Second, the amygdala plays a central role in associative learning processes that assign and update the emotional and motivational value of both rewarding and aversive stimuli [31–33]. Accordingly, the amygdala plays a critical role in alcohol motivation, consumption, and seeking induced by alcohol-predicting cues [33–38]. Importantly, exposure to alcohol cues evokes alcohol craving, which has been correlated with fluctuations of E2 levels [9, 10], and is a major cause of alcohol relapse and use in patients with AUD [39].

Female mice behavior was analysed in the automated IntelliCage system in tests resembling human DSM-5 criteria for AUD [35, 40–42] including: *high motivation to drink alcohol* - measured in a progressive-ratio schedule of reinforcement (motivation test), *high alcohol craving* - measured as alcohol seeking induced by presentation of cues predicting alcohol availability (Cue relapse test), *focus on alcohol use* - measured as alcohol seeking during forced abstinence (extinction test); *persistence of alcohol seeking* - even during signaled alcohol non-availability (persistence test); and *high propensity for relapse* - measured as alcohol consumption upon relapse (alcohol relapse test). Mice were classified as AUD-prone (≥2 criteria positive, top 30% of the population in at least two tests) or AUD-resistant (<2 criteria positive). To identify molecular correlates of vulnerability, we performed RNA sequencing (RNA-seq) of the amygdala, which revealed ERα as a top transcription factor involved in the regulation of differentially expressed genes (DEGs) in alcohol-exposed as compared to alcohol-naive mice. These findings were complemented by the analyses of ERα protein levels in the amygdala, as well as by behavioral and electrophysiological assessments following local genetic manipulation of ERα expression and ovariectomy. Finally, we evaluated associations of *ESR1* single nucleotide polymorphisms (SNPs) with alcohol consumption, craving and loss of control, and blood *ESR1* mRNA levels as a marker and predictor of alcohol use in women with AUD. Overall, our results identify ERα-mediated signaling in the amygdala as an important regulator of cue-induced alcohol seeking, craving, and consumption in females, highlighting the ERα pathway as a potential target for sex-informed therapeutic strategies in AUD.

## MATERIALS AND METHODS

### Subjects

Ten-week old female C57BL/6J mice were obtained from the Nencki Institute animal house. Animals were housed together in standard mouse home cages under a 12/12 hours light/dark cycle. Food and water were available ad libitum. Experimental protocols were designed to minimize the number of animals used and to reduce their suffering. All experiments were conducted in accordance with the Animal Protection Act of Poland guidelines and were approved by the 1st Local Ethical Committee in Warsaw, Poland (no. 823/2019 and 1591/2024).

### AUD model in the IntelliCage

After 1 week of acclimatization, mice were injected subcutaneously (s.c.) with unique microtransponders (8 mm length, 2.12 mm diameter, 0.029 g; Trovan, ID-100) under brief isoflurane anesthesia. Mice were then allowed to recover for 3 days, and the animals with properly located microtransponders were introduced to the IntelliCage system (NewBehavior AG, Zürich, Switzerland) (https://www.tse-systems.com/service/intellicage/), up to 16 animals per system. The IntelliCage consists of a large standard rat cage (20.5 cm high, 40 cm × 58 cm at the top, 55 cm × 37.5 cm at the base). In each corner, a triangular learning chamber is located with two bottles. To drink, only one mouse can go inside a plastic ring (outer ring: 50 mm diameter; inner ring: 30 mm diameter; 20 mm depth into outer ring) that ends with two 13 mm holes (one on the left, one on the right) that provide access to bottle nozzles. Each visit to the corner, nose-poke at the doors governing access to the bottles, and licks were recorded by the system and ascribed to a particular animal.

The training in the IntelliCage was based on published protocol [39] and composed of the following phases: habituation, initiation of alcohol consumption in increasing concentrations, free access to 12% alcohol followed by AUD tests: motivation test, extinction test, cue relapse and alcohol relapse, persistence test, and alternative reward choice. All tests started at the beginning of the dark phase.

#### Habituation phase (day 1-3)

All mice had free access to all bottles with water in both active corners. All doors were open. After 24 hours, when all mice visited and licked from both corners, the doors were closed. Under a fixed ratio of reinforcement (FR 1), each nose-poke was rewarded by a 5-second access to the bottles with water.

#### Free access to alcohol (FA, days 4-47)

During the test, 2 corners were active, each with two bottles available. In one corner, the animals had access to water (“Water corner”), and in the other (“Reward corner”) animals had access to ethanol solution (Alcohol group) at increasing concentrations (4, 8, and 12% ethanol changed every 3-5 days, prepared from 96% ethanol and tap water) or water (Alcohol-naive mice). 12% alcohol was used during all following free access periods (FA2-3). When alcohol (or water) was available, it was signaled by a green cue light turned on in the “reward corner” each time a mouse entered the corner. All liquids were available under an FR1 schedule. 12% alcohol was chosen based on maximal alcohol consumption in g/kg/day during initiation of alcohol consumption. Daily alcohol consumption (g/kg/day) was calculated with the following formula: number of licks of alcohol per day × lick volume [1.94 μl] × alcohol concentration/ animal weight. Our former study showed that the number of alcohol licks calculated by the IntelliCage system correlated with alcohol concentration in the blood [41].

#### Motivation for alcohol tests (days 48-53)

During the test, two corners were active and available to animals. The animals had to perform an increasing number of nosepokes (2, 4, 8, 12, 16, 20, 24, 28, 32, and 36) spaced by less than 1 s during one visit to open the door and be allowed a 5-s access to the reward bottles. The number of required instrumental responses (nosepokes) increased when an animal performed 10 sets of responses of a given ratio. The tests were terminated when 90% of animals did not change the FR level during the last 24 hours. The test usually lasted 7 days. The FR level reached during the test was used as an index of motivation.

#### Extinction of alcohol seeking (days 61-66)

followed by cue (day 67) and alcohol relapse (day 68). The extinction periods were signaled as the “no-reward” periods and lasted 6 days. The door to reward was closed and visits and nosepokes in the reward corner were without scheduled consequences. Average daily number of visits performed in the “reward corner” during the extinction were used as indices of alcohol seeking during forced abstinence. Each extinction was followed by a 24-hour cue-relapse: a green cue light (reward-predicting cue) in the reward corner was presented each time a mouse entered the reward corner. However, nosepokes to the reward doors had no scheduled consequences. A number of nosepokes performed in the “reward corner” during a cue relapse were used as indices of alcohol craving and seeking. This test was followed by alcohol relapse when bottles with alcohol were added to the active “alcohol corner”. During the test each nose-poke into the reward door opened the door for 5 s. The amount of alcohol drank during the first day of relapse (g/kg/day) was used as an index of relapse.

#### Persistence in reward seeking tests

Each persistence test lasted 3 days and was composed of six, 6-hour long “active periods” (A) alternated with 6-hour long “non-active periods” (nA). “Active” periods were signaled by the green cue light in the reward corner. During the “active” periods, nosepokes at all doors opened the door for 5 s (FR 1). The “non-active” periods were signaled by elimination of the green cue light. During the “non-active” periods, nosepokes on the reward side were not followed by any scheduled consequences. The difference of nosepokes performed during nA and A reward periods (Δnosepokes) was used as an index of persistence.

#### Alternative reward test

The test lasted 6 days and during this period mice had unlimited access to 12% alcohol in the alcohol corner (FR 1), and sucrose in increasing concentrations (0.5, 2 and 5% each tested for 2 days) in the other corner. Alcohol (g/kg/day) and sucrose (licks/day) consumption were calculated.

#### Establishment of mouse subpopulations

The addiction index was calculated as previously described [39] and was based on five behaviors: (i) the breakpoint reached during the motivation test, (ii) persistence in alcohol seeking during the persistence test, (iii) alcohol seeking during the cue-induced relapse and (iv) extinction, and (v) alcohol consumption during the alcohol relapse. An individual was considered positive for an AUD-like criterion when its score in the test was in the uppermost 35% of the population. The scoring allowed us to divide the mice into groups according to the number of fulfilled AUD-like criteria: “AUD-prone” who fulfilled 2 or more criteria (≥2 crit. mice); “AUD-resistant” who were positive for one or none of the criteria (<2 crit. mice). Moreover, since the Addiction Index may neglect mice performance in some tests, we also developed Addiction Score. Each of the addiction-like behaviors was normalized and summed up to calculate individual addiction score according to formula: AS = (Vi(individual score) – mean(population))/SD(population). This allowed us to distinguish mice which show consistent behavioral patterns towards alcohol.

### Open-field test (OFT)

Locomotor activity and anxiety-like behavior were assessed using an open-field test. The apparatus consisted of a 63 × 63 cm arena with 32 cm high walls, constructed of grey PVC. The test was conducted in an isolated room under diffuse illumination (70 lux at floor level). Each mouse was placed in the centre of the arena, and its behavior was video-recorded for 20 min. Between mice, the apparatus was cleaned with a 70% ethanol solution (EtOH), wiped with distilled water, and dried. Video data were digitized and analyzed using EthoVision XT15 software (Noldus, Wageningen, NL). Measured parameters included locomotor activity (total distance traveled, mean movement speed, and the number of immobility episodes) and anxiety-like behavior (time spent within the peripheral vs central zones).

### Elevated plus maze (EPM)

Anxiety-like behavior was evaluated using an elevated plus maze (EPM) positioned 50 cm above the floor. The apparatus consisted of two open arms (80 × 5 cm) and two enclosed arms (80 × 5 × 15 cm) extending from a central platform. Each mouse was placed in the centre of the maze and allowed to freely explore all arms for 5 min. Behavioral activity was recorded via a ceiling-mounted camera and analyzed using (EthoVision XT15) software. Between sessions, the maze was cleaned with 70% EtOH and dried. Anxiety levels were quantified based on the time spent in the open arms versus the closed arms.

### Fear Conditioning (FC)

Behavioral experiments were conducted in 31.8 × 25.4 × 34.3 cm chambers (MedAssociates) with a metal-rod floor (8 mm spacing), transparent Plexiglas door, and white walls and ceiling, housed within a soundproof enclosure. A fan provided 0 dB background noise, and near-infrared light was used for illumination.

Two contexts were used. Context A had a metal-rod floor, cleaned with 70% EtOH between animals, with a tray beneath scented with 70% EtOH. In Context B, visual, tactile, and olfactory cues were altered: the floor was covered with grey plastic, a curved white panel with visual cues was added, and the tray was scented with 10% vinegar. Before conditioning, mice were habituated in a separate room for 30 min.

On Day 1, mice were placed in Context A. After 150 s habituation, an auditory cue (10 s) co-terminated with a foot shock (0.7 mA, 2 s) during the final 2 s. This pairing was repeated three times with 30 s inter-trial intervals (ITIs), followed by 30 s in the chamber. On Day 2, mice underwent a cue memory test in Context B: after 150 s habituation, the cue was presented 30 times (30 s ITIs). On Day 3, cue memory was retested in Context B (10 presentations, 30 s ITIs). Six hours later, contextual memory was tested in Context A (30 min, no cue). Sessions were recorded using VideoFreeze software, and freezing was scored manually..

### RNA sequencing

Amygdala tissue was quickly dissected on ice from fresh brain, homogenized and preserved in RNAlater solution (Invitrogen, AM7020) at 4°C for 24 hours, followed by storage at −20°C until further processing. Total RNA extraction was carried out using the RNeasy Mini kit (Qiagen, 74104) in accordance with the manufacturer’s instructions. The RNA concentration, quality, and integrity were assessed using a Nanodrop 1000 (Thermo Scientific) and a Bioanalyzer (Agilent). RNA libraries for sequencing were prepared using the KAPA Stranded mRNA Sample Preparation Kit (KK8401-07962169001, Kapa Biosystems, Wilmington, MA, USA). Sequencing was performed after onboard cluster generation using the HiSeq Rapid SBS Kit v2 (200 cycles) and HiSeq PE Rapid Cluster Kit v2 (Illumina) on a HiSeq 1500 (Illumina). Power, defined as the expected proportion of identified differentially expressed genes among all truly differentially expressed genes, given at least one gene is truly differentially expressed in the data, was modeled using the ssizeRNA R package, assuming that 80% of genes are not differentially expressed, a significance threshold of FDR = 5%, dispersion of 5%, and fold change following a normal distribution. Raw sequencing data were processed using Trimmomatic to remove adapter contamination and eliminate poor-quality reads. Processed reads were mapped to the mm10 reference genome using the STAR algorithm. Subsequently, the Picard MarkDuplicates algorithm was employed to identify and mark optical duplicates. Feature counts from the subread R package were used to quantify the number of fragments assigned to genes. Data normalization and statistical analysis were performed using the NOIseq R package. Quality control analysis of RNA-seq data was conducted using the RSeQC package and STAR (log files). This involved evaluating sequencing fragment distribution, sequencing quality, and the percentage of uniquely mapped fragments. Genes significantly deregulated (FDR-adjusted p-value < 0.05) were used for pathway analysis. Functional enrichment analysis was performed using ShinyGO 0.85.1 [43] to identify affected pathways with a minimum size of 2 genes and maximum 200 genes in the Kyoto Encyclopedia of Genes and Genomes database (KEGG). Transcription factors involved in the regulation of DEGs were identified using the Enrichr platform with the Transcription Factor PPIs database [44–46]. Motif enrichment analysis was performed using Hypergeometric Optimization of Motif EnRichment (HOMER; v3.4). Differentially upregulated and downregulated genes were analyzed using the findMotifs.pl script with the mouse genome annotation. Promoter regions were defined as −1000 bp upstream to +100 bp downstream of the transcription start site (TSS). Enriched motifs were identified using HOMER, and annotated against known transcription factor binding motifs [47]. The Molecular Signatures Database (MSigDB; Mouse MsigDB v2026.1.Mm) Hallmark gene sets were used to identify biological pathways associated with DEGs with a fold change greater than 0.5 and less than −0.5. [48–50].

### Immunostaining

Mice were anesthetized and transcardially perfused with filtered phosphate-buffered saline (PBS, Sigma-Aldrich) followed by 4% paraformaldehyde (PFA, Sigma-Aldrich) in PBS. Brains were then left overnight in the fixing solution and subsequently transferred to 30% sucrose in PBS for 72 hours. Coronal brain sections (40 μm) were obtained using a cryostat (CM1950, Leica Biosystems) and stored at −20°C in an anti-freeze buffer containing PBS, 20% sucrose (Sigma–Aldrich), 15% ethylene glycol (Sigma–Aldrich), and 0.05% NaN_3_ (Sigma–Aldrich). After washing, brain slices were incubated with 5% normal donkey serum (NDS, Jackson ImmunoResearch) in PBS. Slices were incubated with primary antibodies at 4°C (ERα, Millipor (Cat. no. 06-935), overnight, 1:500; PSD-95, Millipore (MAB1598), overnight, 1:500; Arc, Synaptic system (Cat no. 156 003), 72 hrs, 1:1000) in PBS with 0.3% Triton X-100 (PBST) and 5% NDS. The following day, the sections were washed with PBST and incubated with secondary antibodies (Invitrogen, Alexa Fluor 488 (Cat no. A21206), Alexa Fluor 568 (Cat no. A10037)). Subsequently, the slices were washed with PBS, mounted on microscopic slides, and covered with a mounting medium containing DAPI (Fluoromount-G, Invitrogen (Cat no. 00-4959-52)). Immunofluorescent staining was captured using a confocal microscope (Zeiss LSM800, 63 x magnification). All images from one immunostaining were acquired under the same settings. For each animal, at least six representative images were acquired and analyzed using Fiji (ImageJ) [51]. Following smoothing, each channel was converted to 8-bit grayscale. Images were subsequently thresholded for quantification, and mean gray values were measured. All analyses were performed using identical parameters across samples.

### Vaginal lavage and crystal violet staining

To assess the estrous cycle phase of mice, vaginal lavage was conducted according to a previously published protocol [52]. Briefly, the mouse was gently restrained by the tail, and the tip of a sterile 200 μl pipette containing approximately 100 μl of MilliQ water was placed at the opening of the vaginal canal, ensuring not to penetrate the vagina to avoid induction of pseudopregnancy. Subsequently, approximately 50 μl of water was released and aspirated five times. The collected fluid was then spread onto a glass slide and allowed to air-dry at room temperature. Once dried, the vaginal smear was stained with 0.1% crystal violet (Sigma, Cat no. C0775) for 1 minute. Following staining, the slides were rinsed twice in MilliQ water for 1 minute. Excess water was removed, and coverslips were applied to the slides. The estrous cycle phase was then determined under a light microscope. Three primary cell types were identified in vaginal smear samples: well-formed nucleated epithelial cells, cornified epithelial cells, and leukocytes. The relative distribution of these populations varies across the distinct phases of the estrous cycle, enabling stage identification based on the predominant cell type in the smear. During proestrus, smears were dominated by well-formed nucleated epithelial cells. During estrus, cornified epithelial cells predominated and typically exhibited a polygonal morphology. In metestrus, leukocytes became the dominant cell type, although residual cornified epithelial cells could still be present. During diestrus, leukocytes remained abundant, while nucleated epithelial cells began to reappear, indicating the transition to the subsequent proestrus phase [53].

### Electrophysiological ex vivo field recordings

Mice were anesthetized with isoflurane (Iso-Vet, 1000 mg/ml), decapitated, and the brains were rapidly dissected and transferred into ice-cold cutting artificial cerebrospinal fluid (ACSF) consisting of (in mM): 87 NaCl, 2.5 KCl, 1.25 NaH_2_PO_4_, 25 NaHCO_3_, 0.5 CaCl_2_, 7 MgSO_4_, 20 D-glucose, 75 sucrose, equilibrated with carbogen (5% CO_2_/95% O_2_). The brain was cut into two hemispheres and 350 μm thick coronal brain slices were cut using Leica VT1000S vibratome in ice-cold cutting ACSF. The resulting slices were then incubated for 15 min in cutting ACSF at 32°C, followed by minimum 60 min incubation at room temperature in recording ACSF containing (in mM): 125 NaCl, 2.5 KCl, 1.25 NaH_2_PO_4_, 25 NaHCO_3_, 2.5 CaCl_2_, 1.5 MgSO_4_, 20 D-glucose, equilibrated with carbogen.

Extracellular field potential (fEPSP) recordings were conducted in a submerged chamber perfused with recording ACSF in RT. The synaptic potentials were evoked with a custom built stimulus isolator using a concentric bipolar electrode (FHC, 30200) placed in the axons from the basolateral amygdala. The stimulating pulses were delivered at 0.033 Hz and the pulse duration was 0.3 ms. Recording electrodes (resistance 1-4 MΩ) were pulled from borosilicate glass (WPI, 1B120F-4) with a micropipette puller (NARISHIGE, PP-830) and filled with recording ACSF. The recording electrodes were placed in the CeM. Recordings were acquired with MultiClamp 700B amplifier (Molecular Devices, California, USA), digitized with Digidata 1550B (Molecular Devices, California, USA) and recorded with Clampex 10.7 software (Molecular Devices, California, USA). To measure post-synaptic responses the input/output (I/O) curves were obtained by increasing stimulation intensity by 50 μA in the range of 0-700 μA. To measure pre-synaptic release the paired-pulse ratio (PPR) was calculated - two electric stimuli triggering presynaptic action potentials were paired with increasing interstimulus intervals (25, 50, 100, 200 ms) and simultaneously fEPSPs were recorded for each interstimulus interval. Amplitude of fEPSPs was measured using Clampfit 11.1 software (Molecular Devices, California, USA). For the analysis of recordings, we averaged the data from 2 or 3 slices per one animal, 5 mice per group.

### Stereotactic surgeries and virus injections

The surgeries were conducted at least 14 days prior to the beginning of the training. Mice were anesthetized with isoflurane (5% for induction, 1.5–2.0% for maintenance of general anesthesia). Then mice were fixed in the stereotactic frame (51503, Stoelting, Wood Dale, IL, USA). Lentivirus (LV) solution coding short hairpin RNA (shRNA) silencing *Esr1* expression (shEsr1: 500 nl, 8.63 × 10^8^ GC/µl, plasmid: pLKO.1-CMV-tGFP-U6-shEsr1, Sigma, Clone ID: TRCN0000026214; 5’-GCCGAAATGAAATGGGTGCTT-3’) or a control vector based on a pSUPER shRNA targeting the *Renilla* luciferase (5′-CTGACGCGGAATACTTCGA-3′) cloned into pTRIP (shLuc; 500 nl/site; viral titer, 6.52 × 108/µl) were injected into the basolateral amygdala (Bregma AP −0.12, ML +/− 0.24, DV −0.45) according to Paxinos and Franklin [54], at a rate of 0.1 µl per minute, utilizing a nano-injector with glass capillaries (0.58 × 1.00 mm, 100 mm length; Science Products) and 10 µl microsyringe (SGE010RNS, WPI, USA). The capillary was left in place for an additional 10 minutes following injection to prevent outflow. LVs were generated at the Animal Models Laboratory at Nencki Institute (Warsaw, PL).

### Ovariectomy

Ten-week-old C57BL/6J female mice were anesthetized with 5% isoflurane for induction and maintained at 1.5–2% during surgery. Prior to surgery, local antibiotics were administered subcutaneously. Mice were placed in sternal recumbency, and the hip area was shaved and cleaned with 70% alcohol. A 1 cm mid-dorsal incision was made, and the skin was separated from the underlying muscles. Ovaries were identified as white structures beneath the muscles, surrounded by fat pads, and were carefully removed from the uterine horns. Bleeding, if present, was controlled using hemostatic gauze. The uterine horns were returned to the abdominal cavity, and muscles and skin were sutured. Animals recovered on a warm pad. Sham controls underwent the same procedure without ovary removal. Mice were left undisturbed for at least one week before behavioral testing or tissue collection.

### Statistical analysis

Statistical analyses were conducted using GraphPad Prism 11.5 Software. The sample size for each experiment is provided within the figures or their legends. Data with normal distribution and equal variance were subjected to Student’s t-test, one-way ANOVA, two-way ANOVA, followed by *post hoc* Sidak’s tests for multiple comparisons. Least significant difference (LSD) post hoc tests were exclusively utilized for planned comparisons. Non-normally distributed data were analyzed using the Mann-Whitney test for comparison between two groups or the Kruskal-Wallis test for comparison among more than two groups. Spearman correlation was employed for correlation analyses. Data are presented as the mean ± standard error of the mean (SEM). Statistical significance was considered at p < 0.05.

### Clinical genetic ecological momentary assessment study

We analyzed genetic and ecological momentary assessment (EMA) data from the female individuals of the TRR 265 cohort [55–57]. The data were collected between March 2020 and March 2023. The EMA methods are described in Hoffmann et al. [55].

For genotyping, DNA was extracted from blood and/or salivary samples using standard procedures. Genotyping was performed at Life&Brain GmbH, Bonn, using Illumina GSA v3 Arrays.

Genotypes from a total of 754 individuals underwent quality filtering and imputation using the Ricopili pipeline. Quality control (QC) parameters for retaining SNPs and subjects included: SNP call rate ≥ 0.95 (before sample removal) and ≥ 0.98 (after subject removal), subject call rate ≥ 0.98, deviation of autosomal heterozygosity from the mean (|Fhet|≤0.2), minor allele frequency (MAF) ≥ 0.01, and deviation from Hardy-Weinberg equilibrium (HWE p-value > 1e-6). Samples with discordant sex were excluded based on the X chromosome inbreeding coefficient (F), specifically removing pedigree males with Fhet < 0.5 and females with Fhet > 0.5. A SNP was also required to have a valid association p-value.

Principal Component Analysis (PCA) was performed to identify and exclude population outliers, using 92,503 autosomal SNPs with low missingness (<1%) and MAF > 0.05. SNPs were pruned to approximate linkage disequilibrium (LD) using two iterations of LD pruning (r² < 0.2, 200-SNP windows). Known long-range LD regions (e.g., MHC and the chromosome 8 inversion) were excluded. Identity-by-descent (IBD) testing was conducted to identify related pairs using PLINK v1.9. [58]; individuals with PIHAT > 0.2 were identified, and one individual from each pair was randomly removed, prioritizing the retention of cases over controls. The first four principal components (PC1–PC4) derived from this analysis were retained and subsequently included as covariates in all statistical models to account for residual population stratification.

In summary, eight individuals were excluded from the analysis: 4 due to sex errors (i.e., discrepancies between reported and predicted sexes) and 4 due to poor genotype call rates. In total, 36 individuals were excluded as population outliers. One random sample of 30 overlapping pairs were excluded, and 3 related samples were identified.

Finally, genotype imputation was conducted for 680 QC’ed samples using a stepwise pre-phasing and imputation approach, implemented in Eagle2 [59] and Minimac3 [60]. This process utilized default parameters and a variable chunk size of 132 genomic chunks. The imputation reference was the Haplotype Reference Consortium (HRC) panel, which comprises 54,330 phased haplotypes and 36,678,860 variants. Best-guess genotypes were determined by calling genotypes with a posterior probability greater than 0.8; calls not meeting this threshold were set to missing. Moreover, SNPs with missingness > 0.02 were also excluded. Analysis was based on best-guess-genotypes hard called from allele-dosage-data following post imputation QC.

Overall, the female sample with genetic information and all the three alcohol-use phenotypes available comprised 201 participants (N = 200). We report the genotype distributions for the four *ESR1* SNPs (N = 200). For rs6902771, 62 participants (31.0%) carried the CC genotype, 99 (49.5%) the CT genotype, and 39 (19.5%) the TT genotype. For rs11155819, 80 participants (40.0%) carried the TT genotype, 97 (48.5%) the TC genotype, and 21 (10.5%) the CC genotype. For rs6557171, 93 participants (46.5%) carried the CC genotype, 88 (44.0%) the CT genotype, and 19 (9.5%) the TT genotype. For rs2982712, 61 participants (30.5%) carried the TT genotype, 93 (46.5%) the TC genotype, and 46 (23.0%) the CC genotype.

On average, the participants were 38.10 years old (SD = 13.35) with a mean BMI of 24.48 (SD = 4.29; N = 192). Mean body weight was 69.78 kg (SD = 12.95; N = 192) and mean height was 168.77 cm (SD = 6.09; N = 192). They fulfilled on average 4.25 AUD criteria (SD = 1.64), with a maximum lifetime AUD criteria count of 4.48 (SD = 2.26). Of the total sample, 32.3% reported to have a child and 83.3% were in an employed position (N = 192). Regarding daily alcohol use outcomes, participants reported binge drinking on 23.32% of days (SD = 21.17%) and drinking on 54.70% of diary days on average (SD = 25.84%). Mean daily craving was 2.33 (SD = 0.98; N = 195; scale 1 to 7) and mean loss of control over alcohol use was 1.82 (SD = 0.84; N = 184; scale 1 to 6). Mean daily alcohol consumption was 26.52 g (SD = 17.16).

## Statistical Analysis

Generalized linear mixed models (logistic models) were fitted for “binge drinking day” and “day with any alcohol use” as binary dependent variables (yes vs. no) with *ESR1* genotype (rs6902771, rs11155819, rs6557171, rs2982712) as the predictor of interest, using a binomial distribution and a logit link function. Linear mixed models were used for the continuous outcomes craving and loss of control over alcohol use. Each SNP was coded additively according to the number of effect alleles carried: rs6902771 as 0 (CC), 1 (CT), or 2 (TT); rs11155819 as 0 (TT), 1 (TC), or 2 (CC); rs6557171 as 0 (CC), 1 (CT), or 2 (TT); and rs2982712 as 0 (TT), 1 (TC), or 2 (CC).

We report F tests for main effects in all mixed models, followed by t tests for the parameter estimates β. The number of AUD criteria fulfilled, weekend days versus weekdays, COVID-19-related lockdown phase versus no lockdown phase, and the first four ancestry principal components (PC1–PC4) were included as covariates in all models. The AUD criteria count, day type, and lockdown variables have been associated with alcohol use; PC1–PC4 were included to control for residual population stratification, following standard practice in genetic association analyses. To examine whether associations between *ESR1* genotypes and AUD phenotypes changed over the study period, we incorporated genotype-by-time interaction effects (genotype × study day) into the models. The multilevel models included random intercepts to account for the nesting structure of repeated daily observations within participants (level 1: e-diary ratings; level 2: participants). Data were analyzed with SPSS for Windows, version 30.0. Alpha was set at 0.05 (two-sided), without correction for multiple testing.

### Clinical blood *ESR1* mRNA assessment study

A female cohort of 80 early-abstinent alcohol-dependent female patients seeking withdrawal treatment was recruited for the bi-centric, cross-sectional, and prospective Neurobiology of Alcoholism (NOAH) study [61] in parallel to 87 age-matched healthy controls following a multi-step screening procedure in 2013–2014. The Alcohol Use Disorders Identification Test (AUDIT) score was evaluated for the control participants, while the Penn Alcohol Craving Scale (PACS) score as well as lifetime drinking (kg) and daily ethanol intake (kg) were evaluated for the alcohol-dependent patients [61, 62]. The number of readmissions and the days to first readmission were extracted from the electronic patients’ records at the two study centers during a 24-month observation period. Blood samples were obtained from all individuals in the morning following an overnight fast at inclusion (up to 72h of abstinence) and at follow-up (at median 5 days later) into PAXgene TM RNA tubes (Qiagen, Hilden, Germany) and stored at −80°C. RNA was extracted from whole blood using the PAXgene™ Blood RNA System Kit (Qiagen, 762164) employing the manufacturer’s guidelines. RNA concentration and purity were controlled using a NanoDrop ND-1000 UV-visible spectrophotometer (Peqlab, Erlangen, Germany), and the integrity of representative samples was verified on agarose gels in adherence with accepted guidelines for quality control (Taylor et al., 2010). cDNA was synthesized using the High-Capacity cDNA Reverse Transcription Kit (Applied Biosystems™, 4368814). The expression levels of *ESR1* mRNA as well as two reference genes *GAPDH* and *ACTB* (β-actin) were assessed via quantitative PCR (qPCR) using the LightCycler System (LightCycler® SW 1.5, Roche Diagnostics GmbH, Mannheim, Germany). Duplicate reactions were set up in 384-well plates using TaqMan™ Fast Advanced Master Mix (Qiagen, 4444963) and TaqMan Gene Expression Assays Hs01046816_m1, Hs02786624_g1, and Hs99999903_m1 (ThermoFisher Scientific). The geometric mean of the duplicates was adjusted, and the target gene expression was normalized to the geometric mean of the two reference genes [63].

## Statistical Analysis

The data were analysed using IBM SPSS Statistics Version 29 for Windows (SPSSInc., USA) and were visualized using SigmaPlot 15.1.1.26 (Inpixon HQ, USA). Group differences were tested using the Bonferroni-corrected Mann–Whitney U test (3 groups) as the data did not follow a normal distribution in accordance with the Kolmogorov–Smirnov test. The differences between inclusion and follow-up were assessed using the Wilcoxon signed-rank test for related samples. Bivariate correlations were evaluated using Spearman’s method. Subjects with missing data points were excluded from specific analyses. A significance level of p < 0.05 for two-sided tests was considered as statistically significant.

## RESULTS

### ESR1 is the top transcription factor for transcripts deregulated in the amygdala by alcohol drinking

To determine whether estrogen signaling and ERα are associated with vulnerability to develop behaviours characterising alcohol use disorder (AUD) in mouse model, female mice (n = 12) were trained in an IntelliCage-based multidimensional AUD paradigm [35, 40–42] that was composed of 5 tests measuring AUD-related behaviors, including: high alcohol motivation, poor extinction of alcohol seeking during forced abstinence, alcohol craving measured as alcohol seeking during cue relapse, poor control over alcohol consumption during alcohol relapse, and persistence in alcohol seeking during alcohol non-availability periods (see Materials and methods for details). Next, mice were stratified according to DSM-5–like criteria into AUD-prone (positive (top 30% of the population) in ≥2 criteria, n = 5) and AUD-resistant (positive in <2 criteria, n = 7) groups (**Figure 1A**). Behavioral profiling confirmed that AUD-prone animals displayed significantly higher motivation to drink alcohol, alcohol seeking during forced abstinence (extinction test) and cue relapse, higher alcohol consumption during alcohol relapse and free access periods, and higher persistence in alcohol seeking compared with AUD-resistant mice, contributing to overall higher AUD score in ≥2 criteria group and validating the stratification approach (see Materials and methods) (**Figure 1B**). As a control we used mice that were trained in the IntelliCages but they never had access to alcohol (alcohol-naive, n = 5).

**Figure 1.**
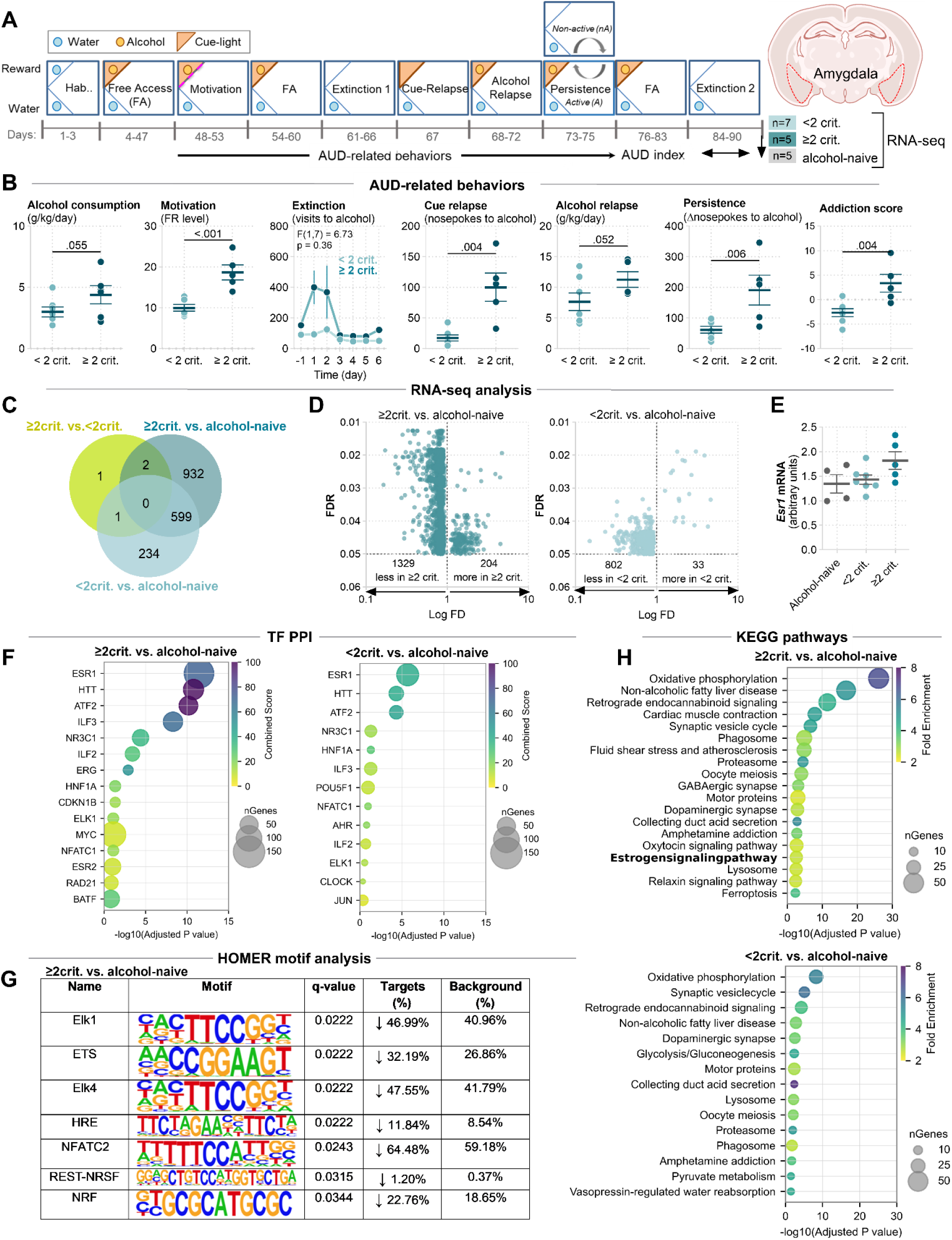
ESR1 is the top transcription factor for differentially expressed genes in the amygdala of mice drinking alcohol. **(A)** Timeline of the IntelliCage-based alcohol use disorder (AUD) paradigm. Female mice underwent alcohol exposure, free access drinking, motivation testing, persistence testing, extinction, cue-induced relapse, and alcohol relapse. Based on DSM-5–like criteria, animals were classified as AUD-prone (≥2 criteria, n = 5) or AUD-resistant (<2 criteria, n = 7). The amygdala tissue was collected after Extinction 2 for molecular RNA sequencing. **(B)** Behavioral phenotyping. Scatter plots show individual performance in AUD-related measures (motivation, persistence, extinction, cue relapse, and alcohol relapse). AUD-prone mice exhibit higher alcohol performance compared with resistant animals (t-tests), confirming successful behavioral stratification. **(C-H)** Transcriptomic analysis. RNA-seq of the amygdala tissue identified differentially expressed genes (DEGs) between ≥2 criteria, <2 criteria, and alcohol-naïve mice. **(C)** Venn diagram illustrates DEG overlap between comparisons, demonstrating a large set of genes uniquely deregulated in AUD-prone mice. **(D)** Volcano plots show magnitude and significance of gene expression changes, revealing stronger transcriptomic remodeling in the ≥2 criteria animals. **(E)** Scatter plot shows *Esr1* mRNA levels quantified in the amygdala. **(F)** Transcription Factor Protein-Protein Interaction (TF PPI) analysis. Plots summarize TFs for DEGs for ≥2 criteria vs alcohol-naïve animals and <2 criteria vs alcohol-naïve animals. **(G)** HOMER motif analysis: Motif discovery reveals overrepresented DNA-binding motifs in regulatory regions of DEGs for ≥2 criteria and alcohol-naïve mice. ↓, downregulated DEGs in ≥ 2 criteria vs alcohol-naïve animals. **(H)** KEGG pathways enrichment analysis. Plots summarize significantly enriched pathways for ≥ 2 criteria vs alcohol-naïve animals and <2 criteria vs alcohol-naïve animals.

Amygdala tissue was collected for RNA sequencing (RNA-seq) after the second extinction test (Extinction 2, day 90) from alcohol-naive (n = 5) and alcohol-trained mice (n = 12) (**Figure 1A**). The comparison of the ≥2 criteria (n = 5 with the highest AUD scores) and alcohol-naive mice yielded 1523 differentially expressed genes (DEGs), including 1329 down- and 204 upregulated in the ≥2 criteria group. Mice with <2 criteria (n = 5 with the lowest AUD scores) had 835 DEGs compared to the alcohol-naive group, 802 down– and 33 upregulated (**Figure 1C**). Nine hundred and thirty two (932) DEGs were unique to the ≥2 criteria group, 234 DEGs were specific for <2 criteria mice and 599 DEGs overlapped between the AUD-prone and resistant animals compared to alcohol-naive mice (**Figure 1D**). Only 4 transcripts significantly differed (at FDR < 0.05) between ≥2 criteria and <2 criteria mice.

Analysis of DEGs from the ≥2 criteria and <2 criteria groups compared with alcohol-naive mice using the Transcription Factor Protein–Protein Interaction library (TF–PPI, Enrichr) [44–46] identified ERα (ESR1) as the top-ranked transcription factor in both comparisons. Notably, the number of ESR1-regulated DEGs was nearly twofold higher in the ≥2 criteria group relative to alcohol-naive mice (132 DEGs) than in the <2 criteria group (73 DEGs) (**Figure 1F; Supplementary Figure 1A; Supplementary Tables 1–2**). Motif analysis of these DEGs using Hypergeometric Optimization of Motif EnRichment (HOMER) revealed enrichment of Elk-1 motif - an ERK-dependent transcription factor downstream of ESR1 signaling and implicated in addiction [64, 65], as well as ETS1 motif, a known ESR1 co-regulator in estrogen response [66], specifically in the ≥2 criteria versus alcohol-naive comparison (**Figure 1G**).

Moreover, pathway enrichment analysis performed using the Kyoto Encyclopedia of Genes and Genomes (KEGG) database identified the estrogen signaling pathway among deregulated pathways in the ≥2 criteria group (16/134 genes, FDR = 0.00059), but not <2 criteria mice (**Figure 1H, Supplementary Table 3-4**), while The Molecular Signatures Database (MSigDB) found *estrogen response: early* and *estrogen response: late* as hallmark pathways in the ≥2 criteria group (**Supplementary Figure 1**). Hence, this data supports the hypothesis that ERα is a key regulator of transcriptomic programs and molecular processes induced in the amygdala by alcohol consumption, and this impact is more pronounced in AUD-prone mice.

Surprisingly, we did not observe significant differences in *Esr1* mRNA levels between the ≥2 criteria, <2 criteria and alcohol-naive groups (**Figure 1E**). However, *Esr1* mRNA abundance (collected on day 90) correlated positively with alcohol seeking during cue relapse (day 67), but not with other AUD behaviors (**Supplementary Figure 2A**). Also, when we compared *Esr1* mRNA levels between the mice stratified according to performance in cue relapse, mice with the highest scores (top 35%) had significantly higher levels of *Esr1* mRNA as compared to the mice with the lowest scores (bottom 35%) (**Supplementary Figure 2B**). Such a difference was not observed when mice were stratified according to scores in other AUD tests. These findings suggest that *Esr1* gene expression in the amygdala is not affected by alcohol consumption, but is a stable trait that is possibly linked to alcohol seeking induced by alcohol-predicting cues.

### ERα protein abundance in the amygdala correlates with alcohol-seeking behavior

To validate mRNA data, we assessed whether ERα protein levels were altered in the mouse amygdala after alcohol training. A new cohort of female mice (n = 27) underwent alcohol training in the IntelliCages, they were tested for AUD-related behaviors, diagnosed as AUD-prone or resistant and sacrificed during free access to alcohol (FA, day 83, n = 14) or after alcohol extinction test (Ext, day 90, n = 13) (**Figure 2A** and **Supplementary Figure 3A-C**). Alcohol-naive mice were used as a control (n = 5). Their brains were sectioned and processed for confocal fluorescent immunodetection of ERα. Quantification of immunofluorescent ERα labeling in the central (CeA) and basolateral (BLA) nuclei of the amygdala showed robust nuclear and neuropile ERα+ signals across both regions (**Figure 2B)**.

**Figure 2.**
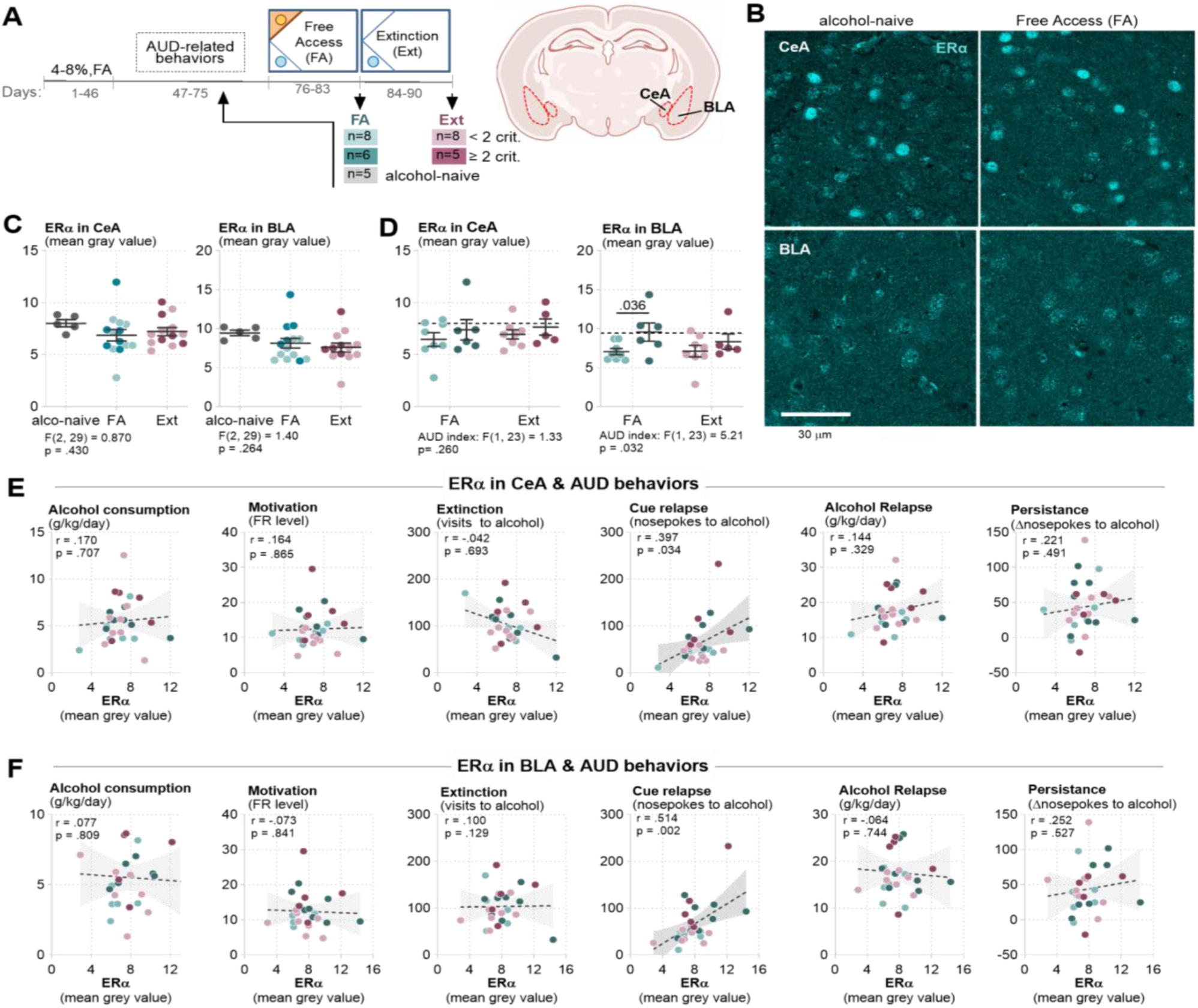
ERα protein levels in the amygdala correlate with alcohol seeking during cue relapse. **(A)** Experimental timeline and sampling. Schematic of the IntelliCage AUD paradigm. Female mice (n = 27) underwent alcohol training in IntelliCages and were phenotyped (<2 criteria - AUD resistant; ≥2 criteria - AUD-prone). Mice were sacrificed after free-access drinking period (FA, n = 14), or extinction (Ext, n = 13) phases. Alcohol-naive mice were used as a control (n = 5). Their brains were collected for histological analyses. Coronal sections of the amygdala were used to quantify estrogen receptor alpha (ERα) protein levels in the central amygdala (CeA) and basolateral amygdala (BLA). The number of animals in each experimental group is indicated in rectangular boxes. **(B)** Representative confocal micrographs showing ERá immunofluorescence in the CeA and BLA. ERá-positive nuclei and neuropile are visible throughout both subregions. **(C-D)** Scatter plots show ERα protein levels quantified in CeA and BLA across behavioral phases and experimental groups. Each dot represents one animal. No significant differences in mean ERα abundance were detected between groups, indicating that overall ERα expression is not altered by alcohol exposure **(C)**, or behavioral phenotype **(D)**. Dashed lines, ERα means for alcohol-naive mice. **(E-F)** Correlation analyses. **(E)** Relationships between ERα protein levels in the CeA and individual AUD-related behaviors. Spearman correlation analyses reveal positive correlations between CeA ERα levels and measures of alcohol cue-induced seeking. Shaded areas represent 95% confidence intervals. **(F)** Analogous analyses and results for the BLA.

Intensity of ERα signal (integrated mean gray value) did not significantly differ between alcohol-exposed and naïve mice neither in CeA nor BLA (**Figure 2C).** We found, however, a slight but significant effect of AUD phenotype on ERα levels in BLA. The ≥2 criteria mice sacrificed during FA had higher levels of ERα as compared to <2 criteria animals sacrificed at the same time (**Figure 2D)**. Moreover, ERα protein levels (in pooled FA and Ext groups as no time effect was found) positively correlated with alcohol seeking during cue relapse both in the CeA and BLA (**Figure 2E-F**), and when mice were stratified according to alcohol seeking during cue relapse the mice with the highest scores had higher ERα signal intensity than mice with the lowest scores (**Supplementary Figure 3D-E**). We found no statistically significant correlations of ERα levels with other AUD behaviors and no difference in ERα signal intensity when mice were stratified based on performance in other AUD tests. These findings indicate that ERα protein levels are not altered by the alcohol training but they correlate with alcohol seeking during cue relapse tested many days prior to the moment when tissue was collected for the molecular analysis.

Importantly, we also found no significant effect of the estrous cycle on ERα levels in the amygdala (**Supplementary Figure 3H**), indicating that ERα levels are relatively stable in the mouse amygdala across estrous cycle. Moreover, estrous phases with high (proestrus and estrus) and low (metestrus and diestrus) estradiol (E2) levels were equally frequent in ≥2 criteria, <2 criteria, and alcohol-naïve mice (**Supplementary Figure 3F-G**), suggesting that neither alcohol training nor AUD phenotype affected estrous cycle in mice.

### Local *Esr1* knockdown in the basolateral amygdala attenuates alcohol seeking induced by cues

To test a causal link between ERα levels in the amygdala and AUD-related behaviors, we used lentiviral vectors encoding shRNAs targeting *Esr1* mRNA (shEsr1) or Renilla *luciferase* as a control (shLuc). shEsr1 and shLuc were injected into the basolateral amygdala (BLA). Two weeks after surgery, allowing for viral expression, mice were trained in IntelliCages to assess general activity and AUD-related behaviors, as in previous experiments. In addition, we included a test measuring alcohol choice over sucrose reward, which reflects reduced valuation of alternative rewards - an established feature of AUD [1] (see Materials and methods for details, **Figure 3A**). *Ex vivo* analysis of the brain sections confirmed LVs expression in BLA. shEsr1 decreased ERα immunoreactivity in transduced cells by ∼75% (**Figure 3B**)

**Figure 3.**
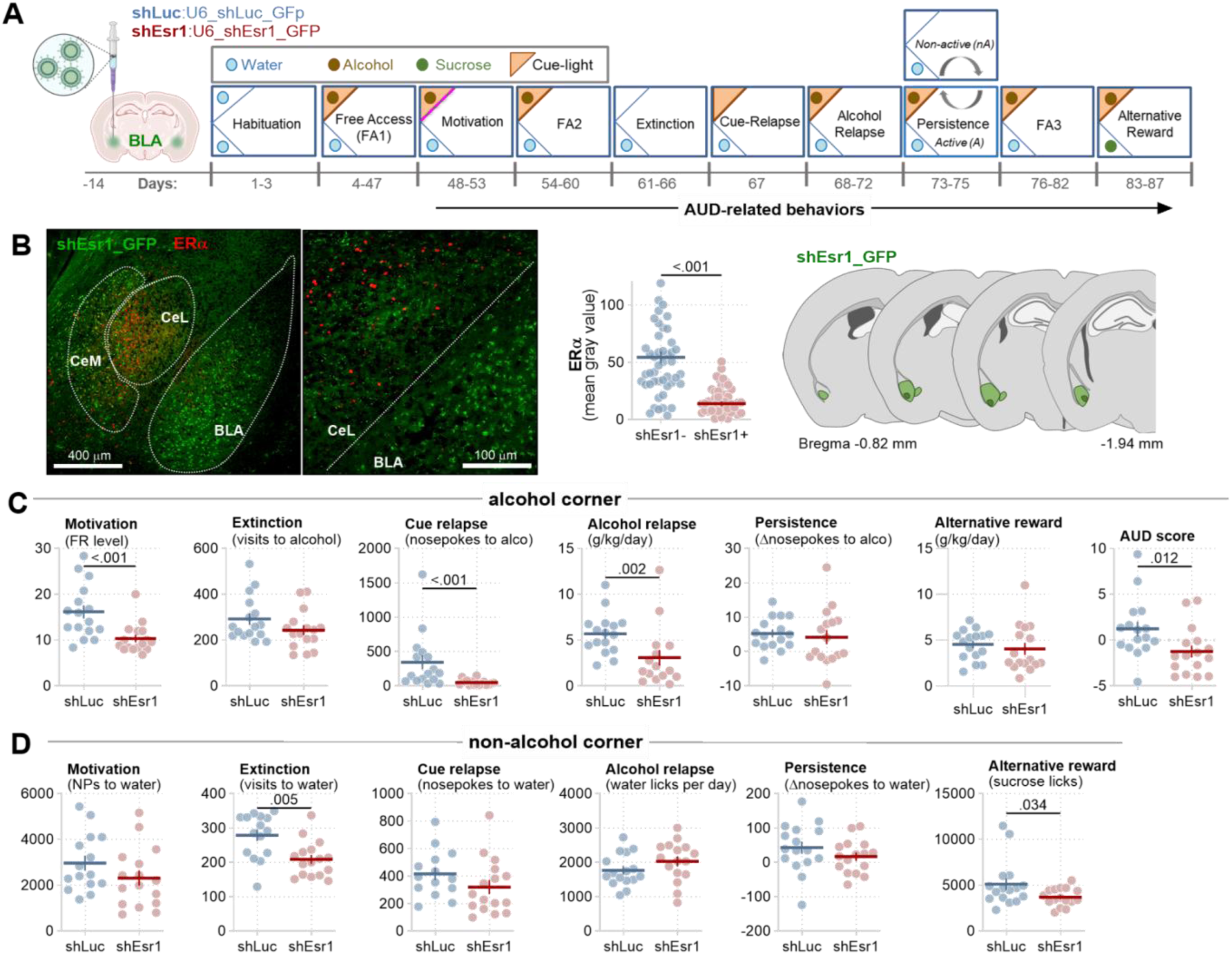
Local knockdown of *Esr1* in the basolateral amygdala attenuates alcohol motivation, cue and alcohol relapse. **(A)** IntelliCage AUD protocol. Mice had shEsr1 (n = 16) or shLuc (n = 16) injected into the basolateral amygdala (BLA). After recovery from surgery and viral expression, mice underwent training in the IntelliCages composed of the following phases: habituation, free access alcohol drinking (FA1-3), and testing of AUD-related behaviors (motivation, extinction, cue relapse, alcohol relapse, persistence and alternative reward test). **(B)** Validation of shEsr1 expression in the amygdala. **(left):** Representative confocal images of the amygdala expressing LVs encoding Esr1 shRNA and GFP (shEsr1) together with ERα immunostaining (red). **(right):** Quantification of ERα immunostaining in shEsr1+ and shEsr1- cells. shEsr1 expression map. pale green, maximal expression; dark green, minimal expression. **(C)** Alcohol-directed measures show decreased alcohol motivation, alcohol seeking during cue relapse and alcohol consumption during relapse, contributing to overall decreased AUD score, in shEsr1 mice relative to shLuc group. shEsr1 has no effect on alcohol seeking during extinction, persistence or alcohol choice over alternative rewards. **(D)** Non-alcohol corner behavior (control). Comparable measures in the non-alcohol (water or sucrose) corner show decreased visits number during extinction and sucrose consumption in the alternative reward tests, but not other behavioral measures, suggesting decreased activity and reward motivation in shEsr1 mice relative to shLuc group. For B-D t-tests were performed, p values when < 0.05 are shown on the graphs.

BLA-targeted *Esr1* knockdown had no effect on mouse activity during the habituation phase and did not affect alcohol consumption during alcohol free access periods, but slightly decreased overall mice activity when they had access to alcohol (total visits, nosepokes) (**Supplementary Figure 4**). More importantly, shEsr1 mice exhibited reduced alcohol motivation, decreased cue-induced seeking, and lower alcohol intake during alcohol relapse, resulting in a reduced overall AUD score (**Figure 3C**). Effects of shEsr1 on mice behavior in the non-alcohol corner were modest and limited to decreased number of visits during extinction, reflecting overall decreased activity, and decreased sucrose consumption, suggesting decreased reward drive (**Figure 3D**). Together, these data demonstrate that ERα signaling within the BLA regulates alcohol motivation and seeking induced by alcohol-predicting cues as well as alcohol consumption during relapse. It also suggests that ERα regulates drive for rewards in general.

Importantly, we did not see the effect of shEsr1 on the frequency of high E2 (proestrus and estrus) and low E2 phases (metestrus and diestrus), suggesting no effect of this local genetic manipulation on E2 levels (**Supplementary Figure 4D**).

### ERα in the amygdala contributes to recall of cue-associated memory

Because alcohol seeking triggered by alcohol-predictive cues requires formation and retrieval of alcohol reward–cue association memory and can be modulated by anxiety - both processes regulated by the amygdala [67, 68] - we examined whether *Esr1* knockdown in the BLA affects anxiety-like behavior and cue memory beyond alcohol cues. To address this, a new cohort of mice (n = 8 per group) underwent stereotaxic surgeries followed by behavioral testing. Anxiety-like behavior was assessed using the open field (OF) and elevated plus maze (EPM) tests, and cue memory formation was evaluated using cued Pavlovian fear conditioning (**Figure 4A**).

**Figure 4.**
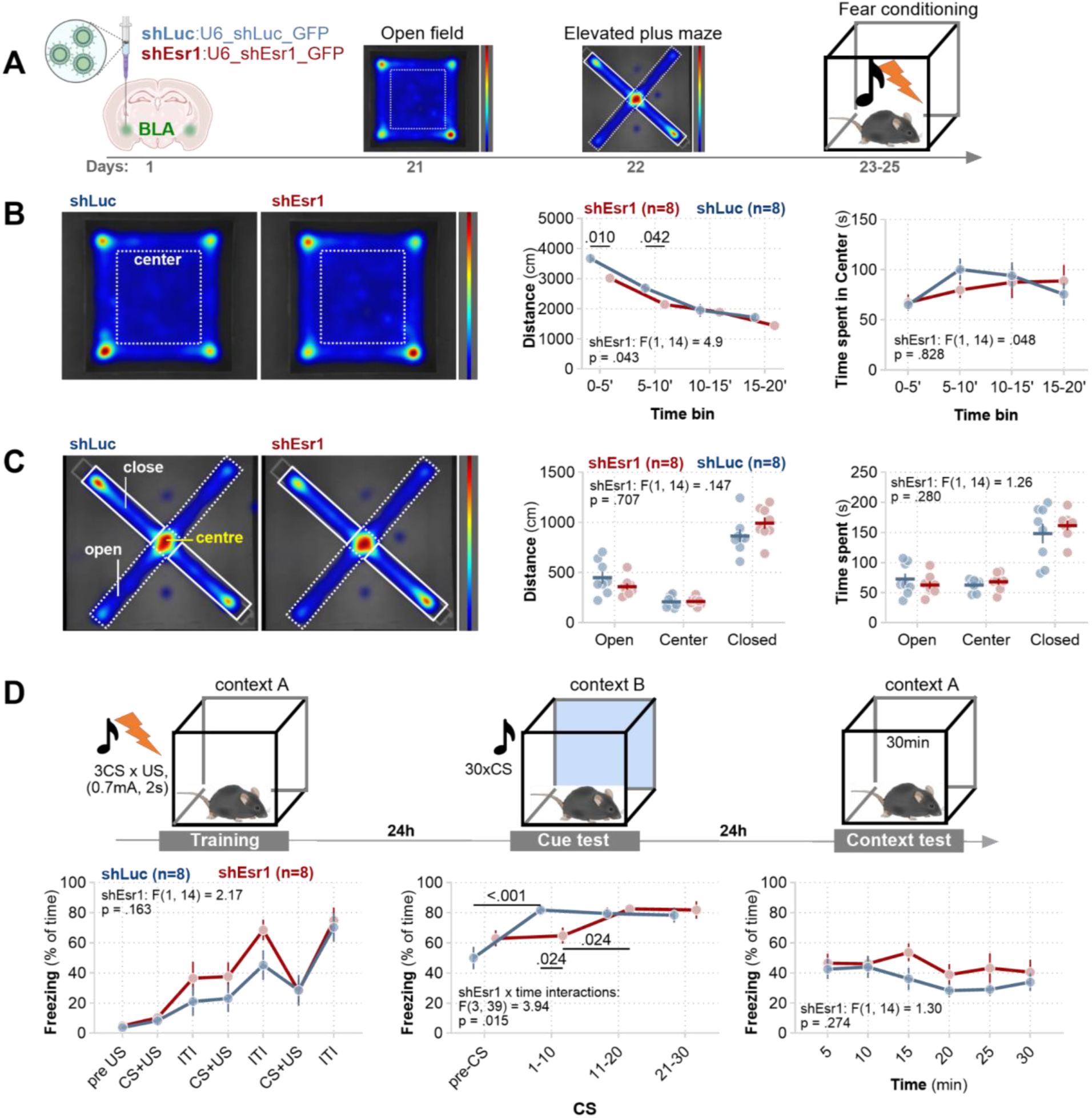
Local knockdown of *Esr1* in the basolateral amygdala alters cue memory recall. **(A)** Experimental timeline. Mice received stereotaxic injections of shRNA constructs expressing control shLuc-GFP (n = 8) or shEsr1-GFP (n = 8) to knock down *Esr1* in BLA. Following recovery, animals underwent open field test, elevated plus maze and cue fear conditioning, followed by cue and context fear memory tests. **(B) (left)**: Representative open-field arena heat maps illustrating exploratory behavior. The dashed square indicates the center zone. **(right)**: Quantification of locomotor and anxiety-like measures across time (total distance traveled and time in center). No major differences in anxiety-like behavior were observed between groups. A significant effect of shEsr1 on mice locomotion was observed. **(C) (left):** Representative occupancy heat maps from an elevated plus maze (EPM). Warmer colors indicate greater time spent. The intersection (“centre”) and distal arms (“close” and “open”) are indicated. **(right):** Quantification of distance travelled and time spent in each zone. Individual animals are shown as dots with group means ± SEM. *Esr1* knockdown does not affect mice behavior. **(D)** Experimental timeline for cued Pavlovian fear conditioning and graphs showing behavioral performance across training sessions for shLuc (blue) and shEsr1 (red) groups. **(left):** During training, shEsr1 mice had slightly increased freezing responses compared with controls, however, the difference was not significant. **(middle):** During the pre-CS phase of Cue testing, no differences were found between groups. However, shEsr1 later increased freezing responses to CSs compared to the controls, indicating impaired recall of cue memories. **(right):** No group-dependent differences in contextual fear memory were found. Points represent mean ± SEM. For B-D, two-way ANOVA tests were performed. Significant effects of shEsr1 and shEsr1 x time interactions are shown on the graphs.

Knockdown of *Esr1* (shEsr1) did not affect anxiety-like behavior, as measured by time spent in the center of the OF or in the open arms of the EPM (**Figure 4B–C**). However, as in the IntelliCages, shEsr1 mice displayed slightly reduced locomotor activity in the OF.

During cued fear conditioning training (**Figure 4D,** left), shEsr1 had no effect on freezing behavior. Both groups showed increased freezing in a new context (context A) following presentation of the first conditioned stimulus (CS) tone co-terminating with the unconditioned stimulus (US; footshock), with further increases during intertrial intervals (ITIs) across the two additional CS–US pairings. In the cue memory test conducted in a novel context (context B; **Figure 4D**, middle), baseline freezing did not differ between the groups. Upon CS presentation, however, only shLuc mice exhibited an immediate increase in freezing during the first 10 CS presentations. In contrast, shEsr1 mice showed a delayed increase in freezing, becoming evident only during later CS blocks (CS 11–20 and 21–30). This pattern suggests intact CS–US memory formation but impaired retrieval of cue-associated memory. Finally, shEsr1 had no effect on freezing during contextual fear recall in context A (**Figure 4D**, right). Together, these findings indicate that ERα signaling in the BLA does not affect anxiety and is not required for the formation of CS–US associations but contributes to efficient cue memory recall and affects mice activity in a new context.

### *Esr1* knockdown in the basolateral amygdala weakens excitatory synapses

Former studies demonstrated the contribution of ERα to synaptic transmission and plasticity of excitatory synapses [21, 26, 69]. As perception of alcohol-predicting cues induces synaptic plasticity in BLA→CeM synapses [34], here we hypothesized that ERα also regulates synaptic strength in this pathway. To address this hypothesis, alcohol-naive female mice were bilaterally injected with shLuc or shEsr1 into BLA (**Figure 5A**). Field excitatory postsynaptic potentials (fEPSPs) were recorded in CeM to evaluate synaptic function by measuring input-output and paired-pulse ratio (PPR) in acute amygdala slices when axons from BLA were monotonically stimulated by increasing stimuli (**Figure 5B**).

**Figure 5.**
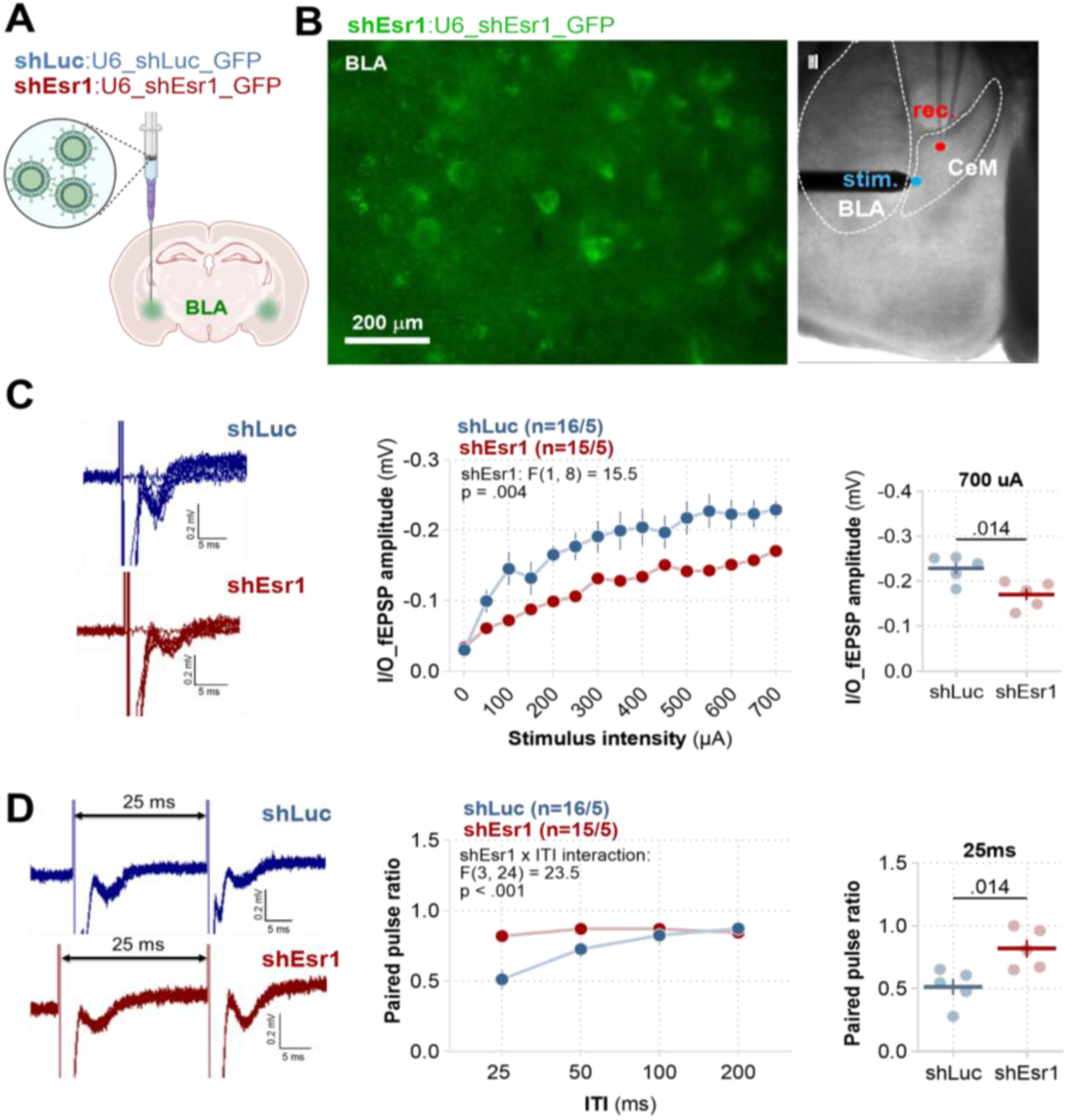
Knockdown of *Esr1* in the basolateral amygdala reduces excitatory synaptic transmission. (A) Schematic of viral injections into the basolateral amygdala (BLA). Mice received bilateral injections of shEsr1 (n = 5) or shLuc (n = 5) into BLA. (B) (**left**): Microphotographs showing shEsr1_GFP-positive cells within the amygdala. (**right**): Amygdala slice microphotography showing recording and stimulation sites. Dashed outlines mark central (CeM) and basolateral amygdala (BLA) boundaries. (C) **(left):** Representative fEPSPs traces evoked by stimuli of different intensities recorded from CeM neurons while the axons from BLA were stimulated. **(middle):** Summary of data for input–output plots of fEPSP amplitude recorded in response to increasing intensities of stimulation. **(right):** Summary of data for fEPSP amplitude elicited by 700 µA stimulus intensity demonstrates reduced post-synaptic responses following Esr1 knockdown, indicating decreased excitatory synaptic strength. (D) **(left):** Example of PPR traces of fEPSPs with 25 ms interstimulus interval. **(middle):** Summary of data for PPR fEPSP amplitude recorded for different inter-stimulus intervals (25, 50, 100, 200 ms). **(right):** Summary of data for PPR fEPSP amplitude recorded for 25 ms interval demonstrates increased PPR responses following Esr1 knockdown, indicating decreased presynaptic release. Numbers of slices/animals per group are indicated in the legends. For C (right) and D (right) t-tests were performed, p values < 0.05 are shown on the graphs. Data are presented as means of a group +/- SEM.

The fEPSP amplitude in the input-output test was significantly decreased in the shEsr1 slices compared to the shLuc (**Figure 5C**). The PPR analyzed as amplitude of fEPSP was significantly increased in the shEsr1 slices, as compared to the shLuc (**Figure 5D**). Thus, *Esr1* knockdown in BLA cells decreased the probability of synaptic release and synaptic transmission in BLA→CeM synapses, suggesting that ERα-regulated synaptic transmission in the BLA→CeM pathway is a plausible mechanism that affects recall of cue memories.

### Ovariectomy reduces ERα signaling in the amygdala and modulates alcohol seeking

To further assess the contribution of estrogen signaling to AUD-related behaviors, we tested how chronic global depletion of E2 by ovariectomy (OVX) affects ERα expression, amygdala activity and AUD phenotype. OVX (n = 18) resulted in the arrest of the estrous cycle (vaginal cytology indicated persistent pseudo-diestrus), and decreased ERα levels in the CeA and BLA as compared to sham surgery (n = 16) (**Supplementary Figure 5A-B**)

We also found that intraperitoneal administration of the ERα agonist, PPT (1 mg/kg), to alcohol-naive mice resulted in lower expression of Arc (an immediate early gene associated with synaptic plasticity [70, 71] and PSD-95 protein (a scaffolding protein and marker of excitatory synapses [72, 73] 90 minutes later in the amygdala of OVX group as compared to the sham animals. Such a difference was not observed between OVX and sham mice treated with saline or E2 (10 µg/kg, i.p.) (**Supplementary Figure 5C**). Hence, OVX chronically decreased ERα expression and ERα signaling linked to plasticity of excitatory synapses.

Next, a new cohort of mice was trained in the IntelliCage system to drink alcohol two weeks after OVX or sham surgery (**Figure 6A**). The OVX group (n = 8) was overall less active compared to the sham controls (n = 7). They made fewer visits and nosepokes in the cage corners, however, total time spent in the cage corners, as well as alcohol consumption during free access, did not differ between the groups (**Supplementary Figure 5**). Because the OVX mice were less active compared to the sham controls even during habituation, we adjusted the assessment of AUD-related behaviors in this cohort to account for reduced activity. Instead of measuring the number of visits and nosepokes, we used time spent in the alcohol-designated corner as an index of motivation, alcohol seeking during extinction, sensitivity to alcohol-predicting cues, and persistence of alcohol seeking (**Figure 6B**).

**Figure 6.**
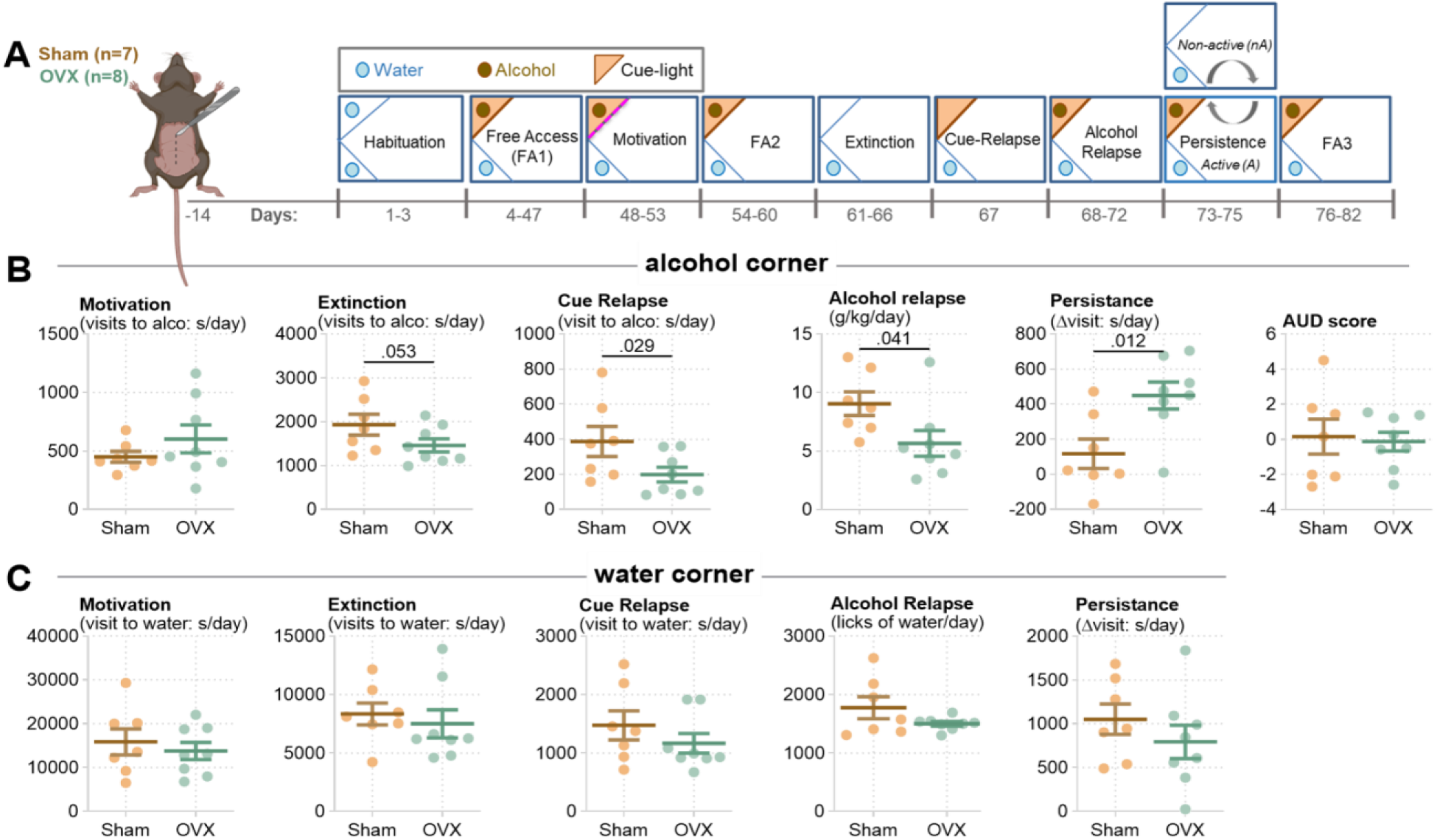
Ovariectomy attenuates cue and alcohol relapse. **(A)** Analysis of AUD behaviors in OVX mice (timeline). After recovery from surgery, sham and OVX mice underwent alcohol training in the IntelliCages composed of: habituation phase, free access alcohol drinking, and testing of AUD-related behaviors including motivation, extinction, cue relapse, alcohol relapse, and persistence. **(B)** Alcohol corner behavior. Scatter plots summarize alcohol-directed measures (e.g., visits, nosepokes, licks/consumption). OVX mice display reduced alcohol-seeking responses during extinction and cue conditions, and increased persistence compared with sham controls. **(C)** Water corner behavior (control). Comparable measures for the water corner show no consistent differences between groups, indicating that behavioral effects are specific to alcohol-related motivation rather than generalized locomotor or exploratory changes. For B and C t-tests were performed, p values when < 0.05, or close to 0.05, are shown on the graphs.

Using these adjusted indices of AUD-related behaviors, we found that OVX mice did not differ from sham controls in motivation for alcohol (**Figure 6B**). However, they showed reduced alcohol seeking during extinction and in response to alcohol-predicting cues, along with lower alcohol consumption during relapse. In contrast, the OVX mice exhibited increased alcohol seeking during the persistence test. No differences between the OVX and sham mice were observed in behaviors associated with the water-designated corner (**Figure 6C**), indicating that the effects were specific to alcohol.

Together, these findings suggest that suppression of estrogen signaling following ovariectomy reduces alcohol seeking in response to alcohol-associated cues and decreases relapse consumption. The results parallel those observed after local BLA *Esr1* knockdown, supporting a model in which ovarian hormones regulate alcohol seeking and relapse consumption via ERα-dependent mechanisms in the amygdala. In contrast, the absence of ovarian hormones appears to impair behavioral flexibility during the persistence test, likely through an amygdala-independent mechanism.

### Human *ESR1* genetic variation is associated with alcohol consumption, craving and loss of control

To translate the preclinical findings into the clinical setting, we analyzed the value of the rs6902771, rs11155819, rs6557171, and rs2982712 *ESR1* gene polymorphisms (**Figure 7A**) to predict real-world daily binge drinking, alcohol use as well as craving for alcohol and loss of control over alcohol intake across the 9-month study period in the TRR265 cohort of women with AUD.

**Figure 7.**
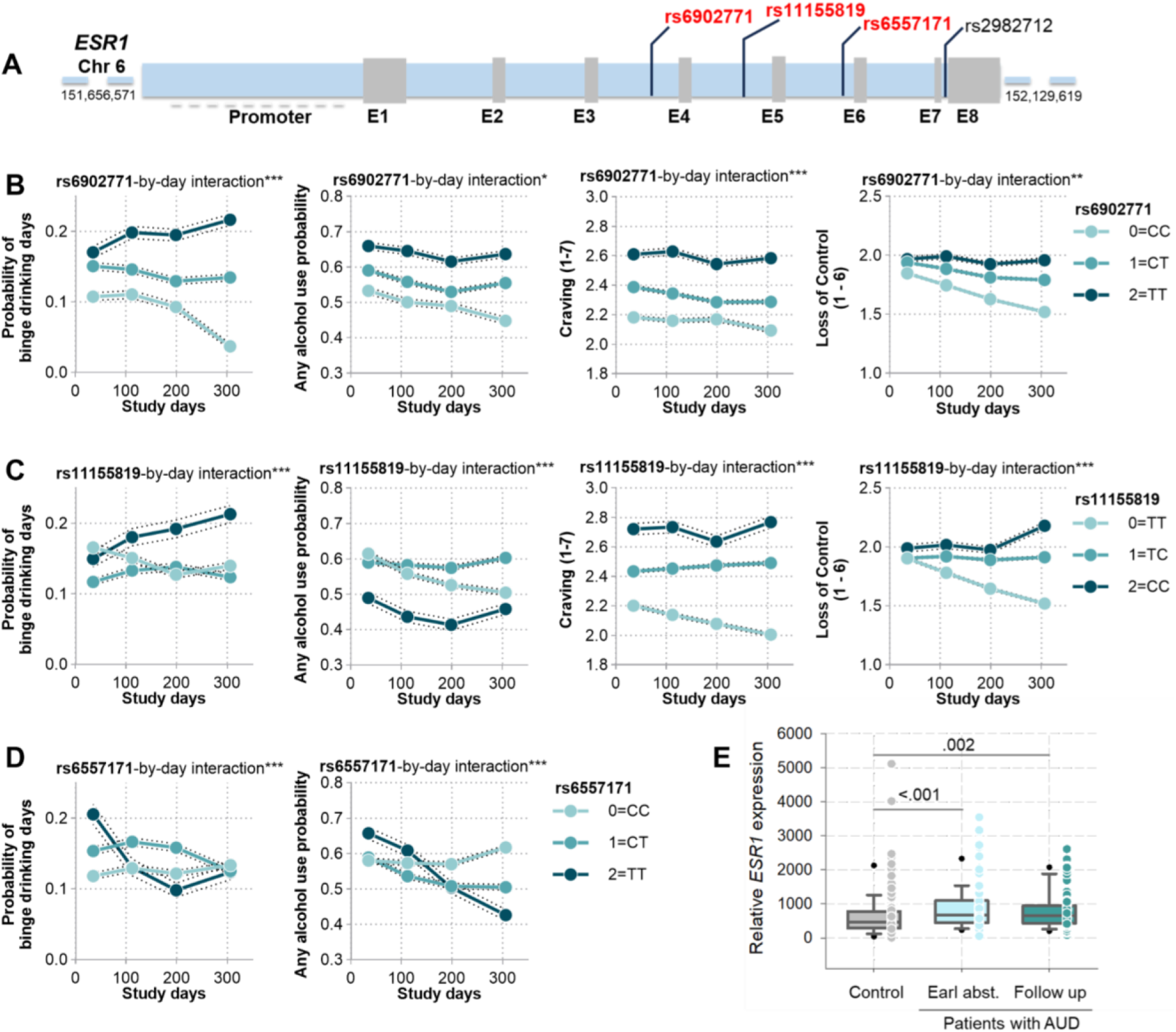
*ESR1* genetic variants and expression patterns in women with AUD. **(A)** Schematic representation of the human ESR1 gene showing the location of analyzed single nucleotide polymorphisms (SNPs): rs6902771, rs11155819, and rs6551717 and rs2982712. Exons are indicated by gray boxes and intronic regions by the light blue line. **(B-D)** Alcohol consumption and craving in the TRR265 cohort of women with AUD divided by genotype for each SNP. To visualize the effects of the SNPs over time, we recoded the continuous day variable into the four study periods of days 1–70, 71–153, 154–244, and 245–367 (= < 25% percentiles, 25% < 50% percentiles, 50% < 75% percentiles, and ≥ 75% percentiles). Within each SNP and day-bin combination, we aggregated the predicted values across participants to obtain mean predicted probabilities and mean craving / loss of control scores and standard errors. (E) *ESR1* mRNA levels are increased in the blood of female patients with AUD from the NOAH cohort at early abstinence and after ca. 5 days follow up in withdrawal treatment (Control: n = 87, AUD patients: n = 80).

We observed a significant rs6902771-by-day interaction effect across all four AUD phenotypes (**Figure 7B** and **Supplementary Table 5**). Higher C-allele dosage was associated with a greater decrease in binge drinking days over time (β = 0.0008, 95% CI [0.0004, 0.0012], OR = 1.0008, p < 0.001) and in days with any alcohol use (β = 0.0003, 95% CI [0.0000, 0.0007], OR = 1.0003, p = 0.044). The same pattern emerged for the continuous outcomes: C-allele dosage was associated with a greater decrease in craving scores over time (β = 0.0005, 95% CI [0.0003, 0.0008], p < 0.001) and in loss of control over alcohol use (β = 0.0006, 95% CI [0.0002, 0.0010], p = 0.003).

Similarly, we found a significant rs11155819-by-day interaction effect on all four AUD phenotypes (**Figure 7C** and **Supplementary Table 6**). Higher T-allele dosage was related to a greater decrease in binge drinking days over time (β = 0.0008, 95% CI [0.0004, 0.0012], OR = 1.0008, p < 0.001) and in days with any alcohol use (β = 0.0011, 95% CI [0.0008, 0.0015], OR = 1.0011, p < 0.001). For the continuous outcomes, higher T-allele dosage was also associated with a greater decrease in craving (β = 0.0005, 95% CI [0.0003, 0.0008], p < 0.001) and loss of control over alcohol use (β = 0.0012, 95% CI [0.0008, 0.0016], p < 0.001).

The rs6557171-by-day interaction effect was significant for the binary but not the continuous AUD outcomes (**Figure 7D** and **Supplementary Table 7**). Higher T-allele dosage was associated with a greater decrease in binge drinking days over time (β = −0.0010, 95% CI [−0.0014, −0.0005], OR = 0.9990, p < 0.001) and in days with any alcohol use (β = −0.0011, 95% CI [−0.0015, −0.0007], OR = 0.9989, p < 0.001). The rs6557171-by-day interaction was not statistically significant for craving (p = 0.172) or loss of control over alcohol use (p = 0.119).

We found no significant rs2982712-by-day interaction effect for any of the four AUD phenotypes: binge drinking days (p = 0.398), days with any alcohol use (p = 0.498), craving (p = 0.902), or loss of control over alcohol use (p = 0.307) (**Supplementary Table 8**).

### *ESR1* blood mRNA levels is increased in woman with AUD diagnosis

To assess predictive and diagnostic value of *ESR1* mRNA blood levels, blood was collected from a cohort of 80 withdrawal seeking early-abstinent female patients with AUD diagnosis from the bi-centric, cross-sectional, and prospective Neurobiology of Alcoholism (NOAH) study and compared to 87 age-matched healthy female controls (**Supplementary table 9**). We observed an increase in the *ESR1* mRNA expression in the patients, both at the hospital admission (p < 0.001) and after 5-day detoxification (p = 0.002; **Figure 7E**). This parameter did not significantly change after detoxification compared to the early abstinence level (p = 0.852), and a strong correlation was observed between these values (r = 0.497, p < 0.001). However, no correlations between *ESR1* mRNA expression and lifetime or daily alcohol drinking, score of Penn Alcohol Craving Scale (PACS), number of readmissions or days to first readmission were found (**Supplementary table 10**). These findings indicate that peripheral *ESR1* mRNA levels can be considered as a marker of AUD, however it does not predict alcohol-related readmissions to hospital.

## DISCUSSION

In this study, we provide convergent evidence from transcriptomic, molecular, circuit-level, behavioral, and human genetic analyses that estrogen receptor alpha (ERα; ESR1) signaling in the amygdala is an important regulator of alcohol-seeking behavior and relapse vulnerability. By combining an ethologically relevant multidimensional AUD model in mice with causal manipulations and human ecological momentary assessment and genetic data, we identify amygdala ERα not merely as a marker, but as a functional driver of cue-induced alcohol seeking.

A key finding of this work is that estrogen signaling emerges as a selectively deregulated pathway in the amygdala of AUD-prone animals. ESR1 was identified as the top transcription factor associated with DEGs in alcohol-exposed mice and ESR1-regulated DNA motifs were specifically enriched for DEGs in vulnerable mice, indicating that ERα-dependent transcriptional programs may underlie susceptibility to maladaptive alcohol-related behaviors. Importantly, neither Esr1 mRNA nor ERα protein levels differed between experimental groups on average, yet both measures positively correlated with cue-induced alcohol seeking. This suggests that inter-individual variability in ERα signaling, rather than gross expression changes induced by alcohol exposure, encodes vulnerability to relapse-like behavior. Such trait-like associations are consistent with stable individual differences in addiction susceptibility [74–76].

Previous work has established that estrogen signaling modulates alcohol-related behaviors in VTA, BNST and dorsal raphe nucleus, where estrogen enhances alcohol-induced neuronal activity and promotes drinking both in male and female mice [26, 27, 77, 78]. Our causal experiments extend previous work showing that ERα signaling in the basolateral amygdala (BLA) is necessary for the expression of key AUD-related behaviors in female mice. Local knockdown of *Esr1* reduced alcohol motivation, cue-induced responses, and relapse drinking, supporting a direct role of ERα in promoting alcohol-directed behavior. These effects were not accompanied by major alterations in anxiety-like behavior, indicating that ERα specifically modulates motivational and associative processes rather than general affective states. However, a modest reduction in locomotor activity and sucrose consumption suggests that ERα in BLA may contribute more broadly to reward sensitivity supporting earlier study [79].

Mechanistically, our electrophysiological data indicate that ERα regulates excitatory synaptic transmission within the amygdala, specifically in the BLA→CeA pathway. *Esr1* knockdown reduced synaptic strength and decreased presynaptic release probability, pointing to ERα as a modulator of glutamatergic transmission. This is consistent with prior evidence that ERα enhances excitatory synaptic transmission and plasticity [22, 80–83]. Given that BLA→CeA projections convey associative information about cue–reward contingency [33–35, 84], ERα-dependent modulation of this synapse provides a direct mechanism for controlling the strength of cue representations. We propose that ERα enhances synaptic efficacy within this pathway, thereby facilitating the propagation of cue-related information to downstream output circuits that drive alcohol seeking.

Consistent with this synaptic mechanism, ERα knockdown selectively impaired the retrieval of cue-associated memory without affecting acquisition. This phenotype suggests that ERα is not required for encoding cue–reward associations but is critical for their effective reactivation. At the circuit level, reduced excitatory transmission in the BLA→CeA pathway likely lowers the signal-to-noise ratio of cue-evoked activity, resulting in delayed or attenuated behavioral responses to conditioned stimuli. This interpretation aligns with the observed reduction in cue-induced alcohol seeking following *Esr1* knockdown.

Hormonal manipulation further supports a role for endogenous estrogen signaling in regulating alcohol-related behaviors. Ovariectomy reduced expression of ERα and synaptic plasticity markers (Arc, PSD-95) in the amygdala, indicating diminished activity-dependent synaptic remodeling, accompanied by decreased cue-induced alcohol seeking and relapse drinking. These effects closely mirror those observed following local *Esr1* knockdown, reinforcing the conclusion that ovarian hormones act, at least in part, through ERα-dependent mechanisms in the amygdala. Together, these findings support a model in which endogenous estrogen signaling maintains ERα-dependent synaptic potentiation within amygdala circuits. Loss of this signaling reduces excitatory drive and weakens the behavioral impact of alcohol-predictive cues. Interestingly, ovariectomy increased persistence in alcohol seeking, suggesting that distinct components of AUD-like behavior may be differentially regulated by estrogen signaling and may involve additional brain regions or compensatory mechanisms.

At the systems level, our findings position ERα as a neuromodulator of amygdala circuit excitability. ERα is known to couple to metabotropic glutamate receptors and activate intracellular signaling cascades that regulate synaptic strength, including MAPK/ERK and CREB-dependent transcription [23, 24, 85–88]. Through these pathways, ERα may dynamically adjust synaptic efficacy and neuronal responsiveness to salient stimuli. In the context of AUD, elevated ERα signaling would be expected to amplify cue-evoked activity within amygdala circuits, thereby increasing craving and relapse risk.

Importantly, our findings translate to humans. *ESR1* polymorphisms associated with higher receptor expression predicted increased alcohol binge and daily consumption as well as craving and loss of control. Importantly, the same variants were linked to familial risk for AUD [29, 89]. This may therefore serve as potential biomarkers for risk stratification and treatment response. Moreover, we observed increased blood levels of *ESR1* mRNA in women with AUD diagnosis. Overall, these data provide strong cross-species validation and suggest that ERα signaling contributes to individual differences in AUD vulnerability in humans, paralleling the trait-like associations observed in mice.

Together, our results support a model in which ERα signaling in the amygdala enhances excitatory synaptic transmission and facilitates the retrieval of alcohol-associated cue memories, thereby promoting alcohol seeking and relapse. This mechanism may be particularly relevant in females, which have higher brain expression of ERα [90] and where fluctuations in estrogen levels dynamically modulate ERα activity, potentially contributing to variability in craving across the menstrual cycle [91].

Several limitations should be acknowledged. First, while our study focuses on the amygdala, ERα is widely expressed across the brain, and its contribution to AUD likely involves additional regions such as the prefrontal cortex and mesolimbic dopamine system. Second, the observed effects of ovariectomy and knockdown on general activity and reward processing indicate that ERα may have broader roles that should be further dissected. Third, although our human genetic data support translational relevance, functional validation of *ESR1* variants in neural circuits activity remains to be established.

Despite these limitations, our findings identify ERα as a key molecular and circuit-level regulator of alcohol-seeking behavior and highlight estrogen signaling as a promising target for sex-informed therapeutic strategies in AUD. Pharmacological modulation of ERα, including the use of selective estrogen receptor degraders or modulators, may offer novel approaches for addressing the “telescoping effect” in women with AUD [6–8].

## AUTHOR CONTRIBUTION

Conceptualization: RP, LK, EC, JS, BL, KR. Investigation: RP, LK, MP, YR, JS, BW, BG. Visualization: RP, LK, KR. Supervision: BL, KR. Writing—original draft: RP, LK, KR. Writing—review & editing: all authors.

## DATA AVALIABILITY

RNAseq data is available at GEO NCBI (<u>GSE221166</u>). The other datasets generated during the current study will be available at Open Science Framework upon acceptance for publication.

---

**Supplementary Figure 1.**
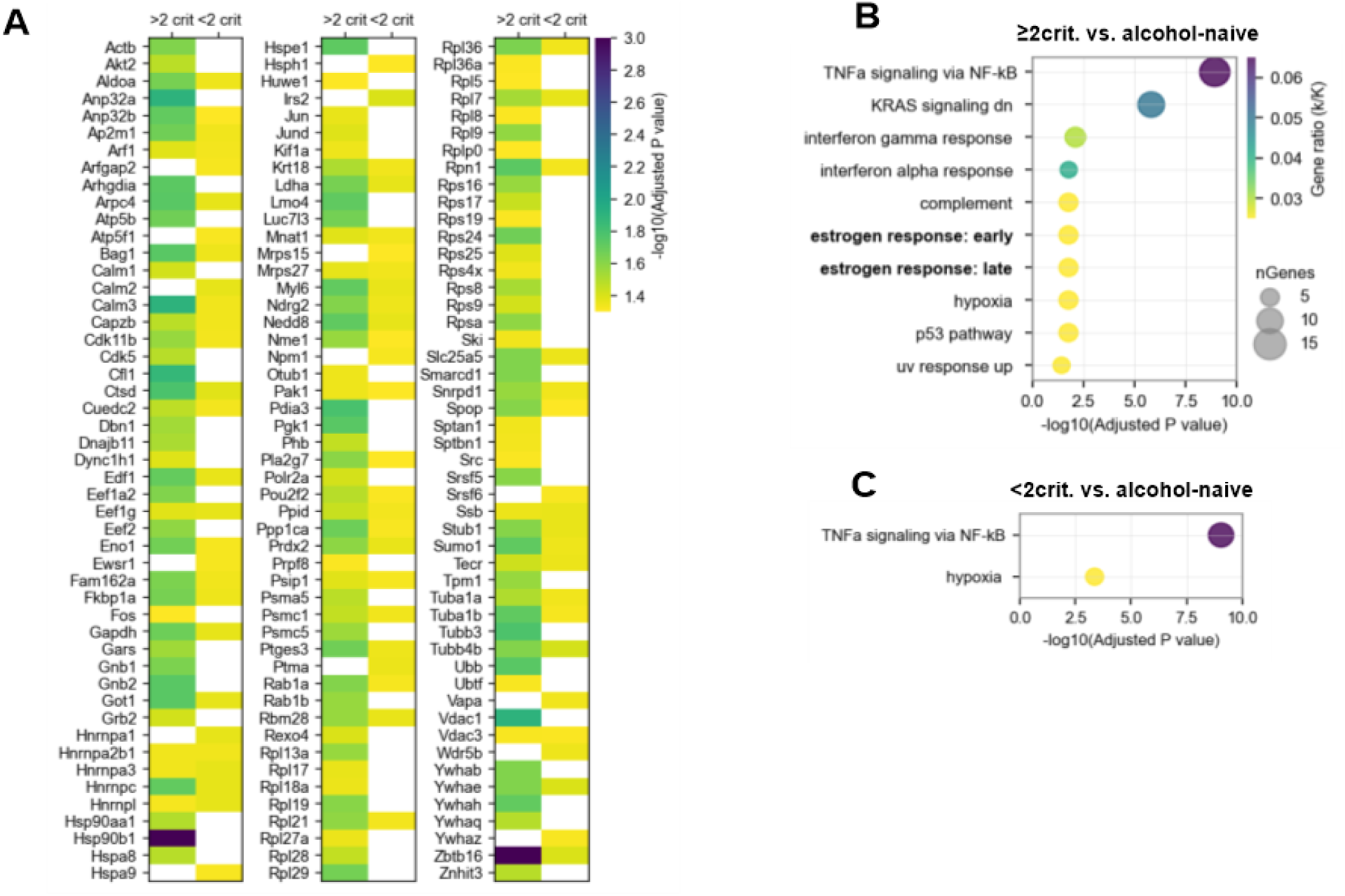
The ESR1-target DEGs and estrogen signalling pathway is selectively deregulated in the amygdala of ≥2 criteria mice. (A) TF PPI analysis. DEGs regulated by ERα (ESR1) in ≥2 and < 2 criteria mice. (B-C) MSigDB hallmark pathways enrichment analysis. Plots summarize significantly enriched pathways for ≥2 criteria vs alcohol-naïve animals (B) and <2 criteria vs alcohol-naïve animals (C). Functional enrichment highlights the estrogen response: early and late among the hallmark pathways in ≥2 criteria mice (Supplementary Table 5).

**Supplementary Figure 2.**
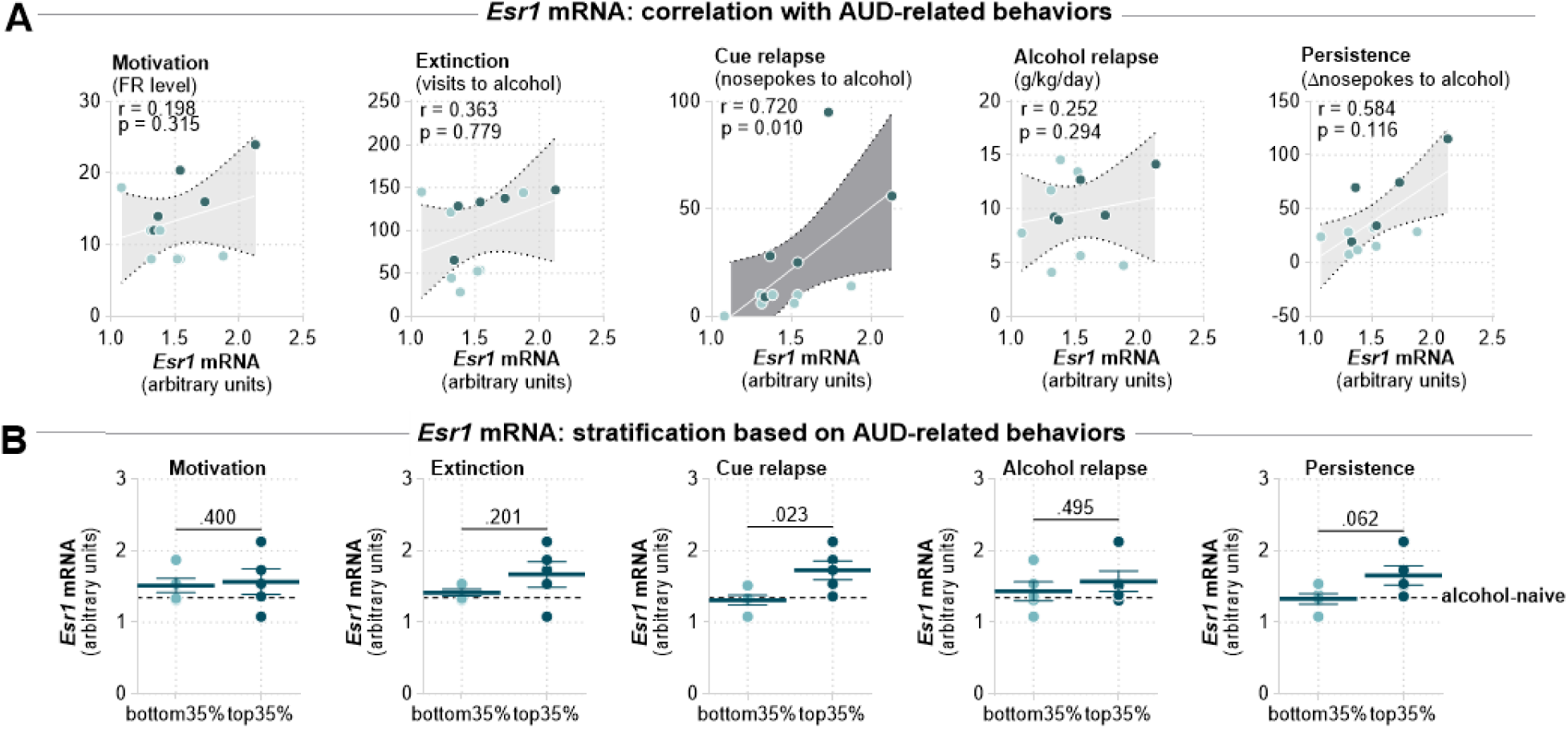
Performance in cue relapse predicts *Esr1* mRNA levels in the amygdala. **(A)** Molecular-behavioral correlations. Correlation analyses demonstrate that *Esr1* mRNA levels in the amygdala positively associate with alcohol-seeking behaviors, particularly during extinction, cue-induced seeking and persistence test, whereas no associations are observed for other behavioral parameters. Each point represents an individual animal; shaded areas indicate confidence intervals. Spearman correlations were calculated. **(B)** Mice were stratified according to performance in AUD tests. bottom 5, mice with the lowest performance in the test; top 5, mice with the highest score in the test. Dashed line, mean *Esr1* mRNA levels in the amygdala of alcohol-naive mice. t-tests were performed, each dot represents individual mouse data, means +/_ SEM are shown. t-tests were calculated.

**Supplementary Figure 3.**
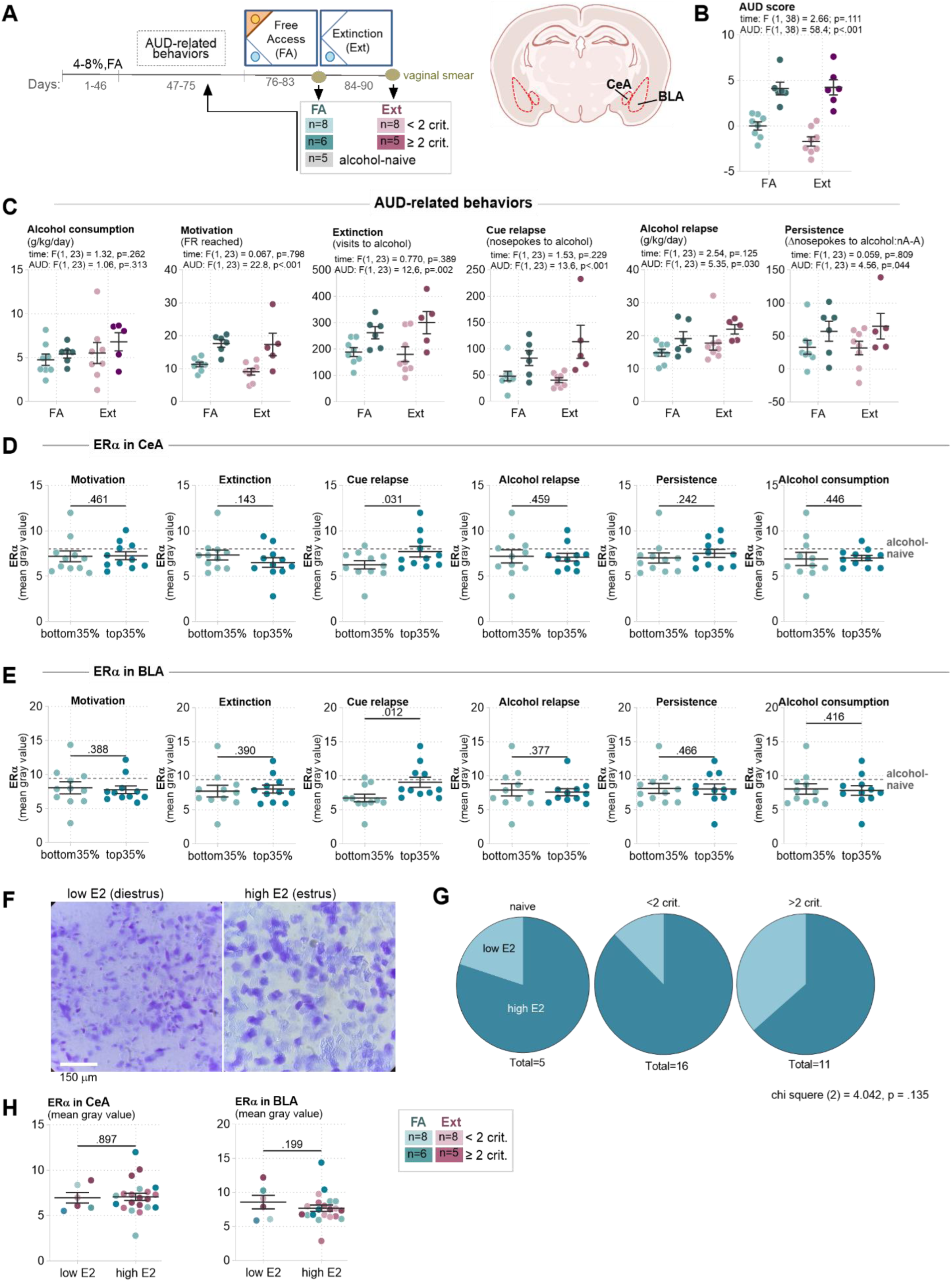
Experimental timeline, AUD-related behaviors and estrous cycle assessment. **(A)** Experimental timeline. Female mice were trained in the IntelliCags to drink alcohol. AUD-related behaviors were assessed to determine AUD index (<2 criteria - AUD resistant; ≥2 criteria - AUD-prone), and animals were sacrificed following free access to alcohol (FA) or extinction (Ext). Alcohol-naive mice were used as a control. Animals were perfused and coronal sections of the amygdala were used to quantify estrogen receptor alpha (ERα) protein levels in the central amygdala (CeA) and basolateral amygdala (BLA). Vaginal smears were collected before perfusion to determine estrous cycle stage. Group sizes are indicated (FA and Ext groups; n values shown in figure). **(B-C)** AUD score and AUD-related behavioral measures plotted according to AUD index (<2 criteria vs ≥2 criteria) and experimental condition (FA vs Ext). Behavioral parameters include alcohol consumption (g/kg/day), motivation (progressive ratio), extinction of seeking during forced abstinence, impaired control measures (cue-induced alcohol seeking), persistence during alcohol non-availability, excessive intake during relapse as defined in the Methods. Individual data points represent single animals. Group means ± SEM are shown. **(D-E)** Mice (n = 27) were stratified according to performance in AUD tests. Mice performance during cue relapse, but not other AUD tests, predicts ERα protein levels in CeA **(D)** and BLA **(E).** Grey dashed lines, alcohol-naive mice; bottom 35%, mice with the lowest performance in the test; top 35%, mice with the highest scores in the test. t-tests were performed, each dot represents individual mouse data, means +/_ SEM are shown. **(F)** Representative images of vaginal cytology used to classify estrous cycle phases corresponding to low (metestrus and diestrus) and high (proestrus and estrus) E2 states. **(G)** Distribution of animals across low and high E2 estrous phases within each experimental group (pie charts). **(H)** Scatter plots show ERα protein levels quantified in CeA and BLA across estrous cycles. Each dot represents one animal. No significant differences in mean ERα abundance were detected between estrous cycle phases. Individual values and group means ± SEM are shown.

**Supplementary Figure 4.**
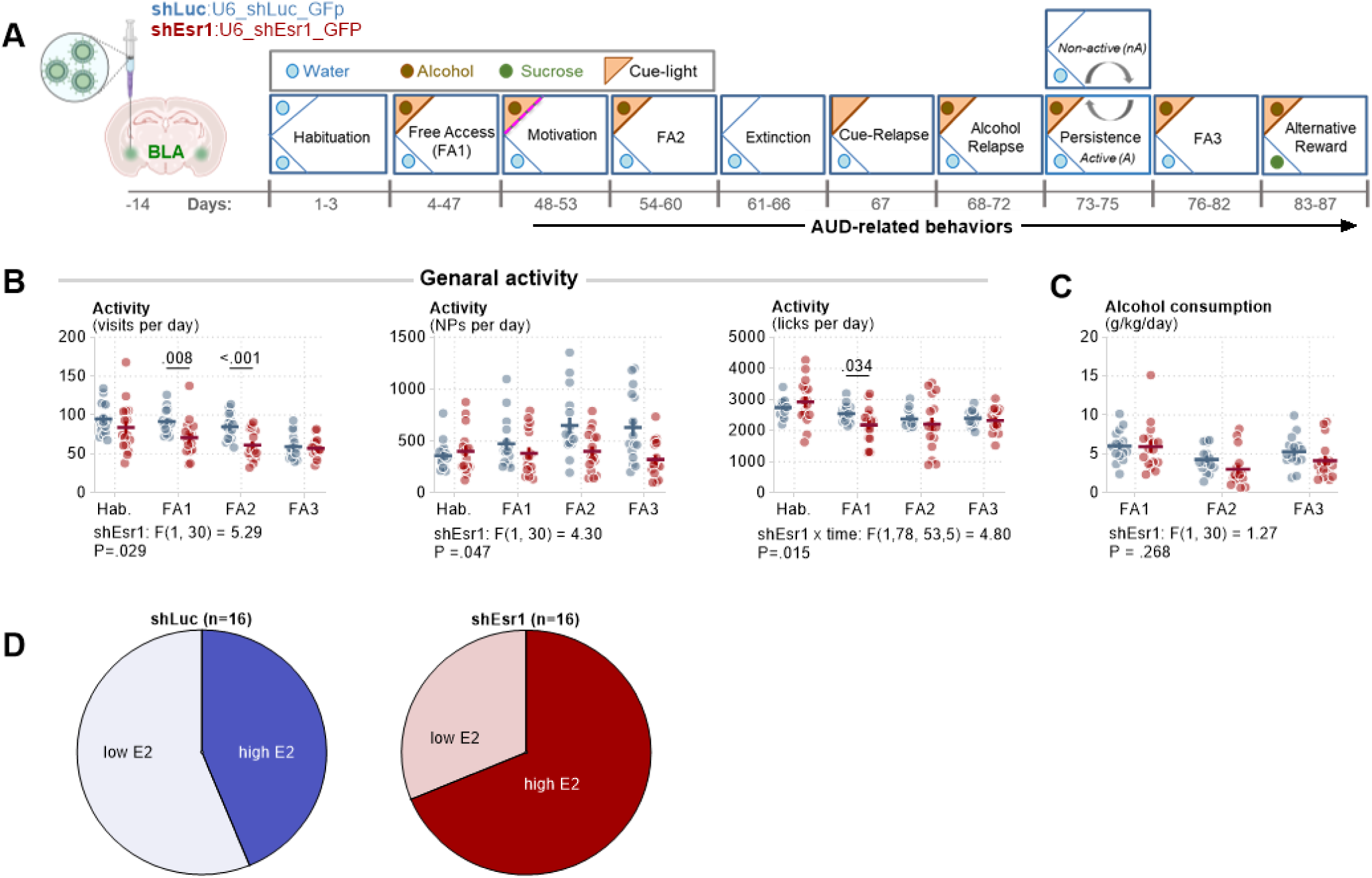
Knockdown of *Esr1* in the basolateral amygdala slightly decreases mice activity but does not affect alcohol consumption or estrous-cycle. (A) Validation of shEsr1 LV efficiency to downregulate ERα protein in neuronal culture. (left): Microphotographs showing shEsr1_GFP-positive cells (green) and ERα fluorescent immunostaining (magenta). (right): Quantification of data showing that shEsr1 downregulated ERα by approx. 40%. (B) IntelliCage AUD protocol. Mice had shEsr1 (n = 16) or shLuc (n = 16) injected into the basolateral amygdala (BLA). After recovery from surgery and viral expression, mice underwent training in the IntelliCages composed of the following phases: habituation, free access alcohol drinking (FA1-3), and testing of AUD-related behaviors (motivation, extinction, cue relapse, alcohol relapse, persistence and alternative reward test). (C) Summary of data comparing shLuc and shEsr1 mice activity. Each point represents one mouse. The group means ± SEM are shown. *Esr1* knockdown decreased general activity in mice drinking alcohol (visits and nosepokes in FA1-3), as well as total licks (water and alcohol). (D) *Esr1* knockdown had no effect on alcohol consumption during alcohol free access phases (FA1-3). (E) Pie charts show comparable estrous-cycle phase distribution between shLuc and shEsr1 groups, indicating that local ERα manipulation does not affect estrous-cycle.

**Supplementary Figure 5.**
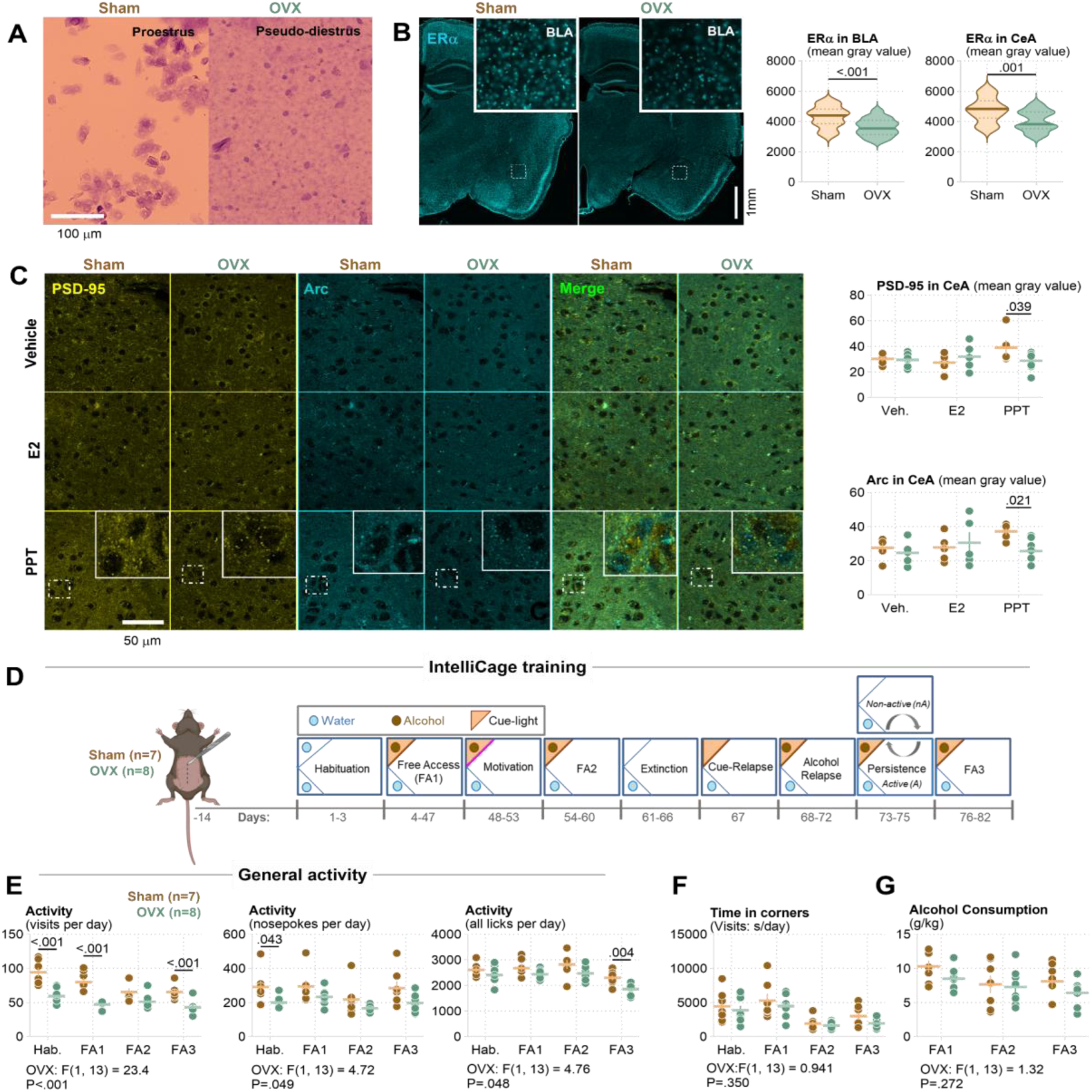
Ovariectomy reduces ERα expression and signaling in the amygdala, and modulates mice activity in the IntelliCages. (A-B) Verification of ovariectomy and ERα expression. 10-week old mice underwent OVX or sham surgery and 2 weeks later they were perfused and their brains sliced for fluorescent immunodetection of ERα. (A) Representative vaginal cytology confirming successful ovariectomy in OVX mice (OVX; pseudo-diestrus–like cytology). (B) Representative coronal sections of the amygdala sections immunostained for ERα protein, with higher-magnification images of the basolateral amygdala (BLA) inset. Right, quantification of ERα immunoreactivity in BLA and CeA showing reduced ERα protein levels in OVX mice compared with sham controls. Mice were sacrificed 2 weeks after OVX. (C) Synaptic protein analysis in CeA of OVX mice. 10-week old mice underwent OVX or sham surgery and 2 weeks later they were injected with E2 (10 µg/kg, i.p.), PPT (1 mg/kg, i.p. or vehicle; n = 5-7 per group). Mice were perfused 90 minutes after the drug injection and their brains sliced for fluorescent immunodetection of Arc and PSD-95. (left): Representative confocal images of PSD-95, Arc and merged immunofluorescence in CeA from Sham and OVX mice. Insets show higher-magnification regions of interest. (right): Plots quantify PSD-95 and Arc signal intensity. OVX have lower PSD-95 and Arc levels as compared to the Sham group after PPT treatment. (D-G) Analysis of AUD behaviors in OVX mice. (D) Behavioral timeline in the IntelliCages. After 2-week recovery from surgery, sham and OVX mice underwent cage habituation, free access alcohol drinking (FA), and testing of AUD-related behaviors including motivation, extinction, cue relapse, alcohol relapse, and persistence. (E) General activity. Plots show mice activity across training phases for sham and OVX animals. OVX animals exhibit reduced activity (visits and nosepokes per day), while acquisition of alcohol corner preference (visits (s) per day) (F) and consumption (alcohol g/kg/day) (G) are largely preserved.

**Supplementary Table 1.**
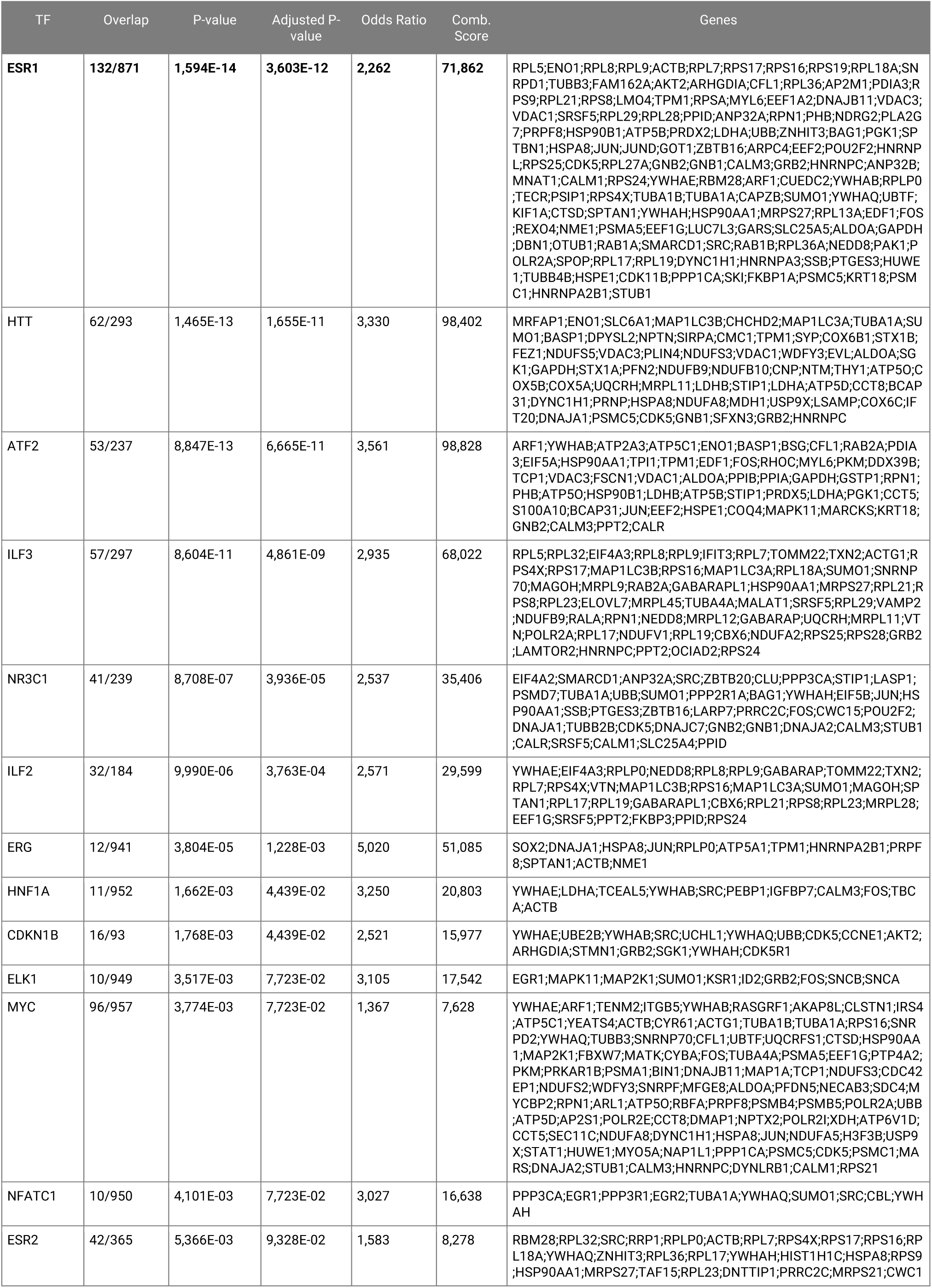
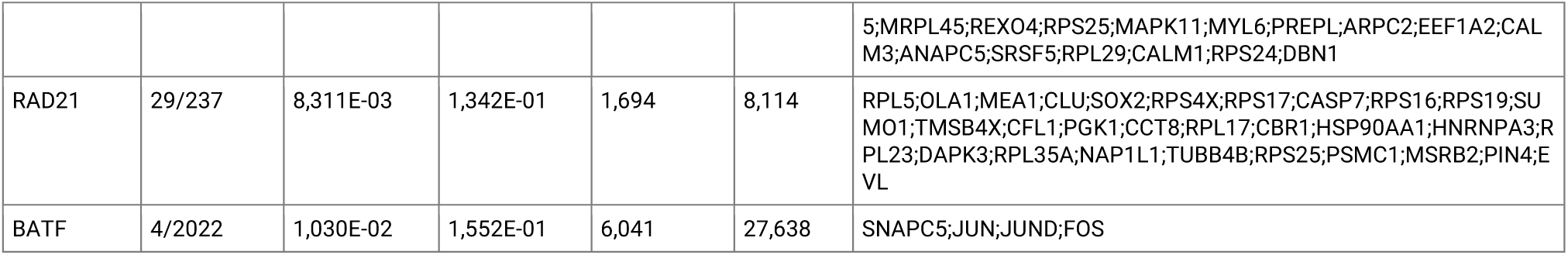
Transcription Factor Protein-Protein Interaction (TF PPI) analysis for the amygdala DEGs for ≥2 criteria vs alcohol-naive mice.

**Supplementary Table 2.**
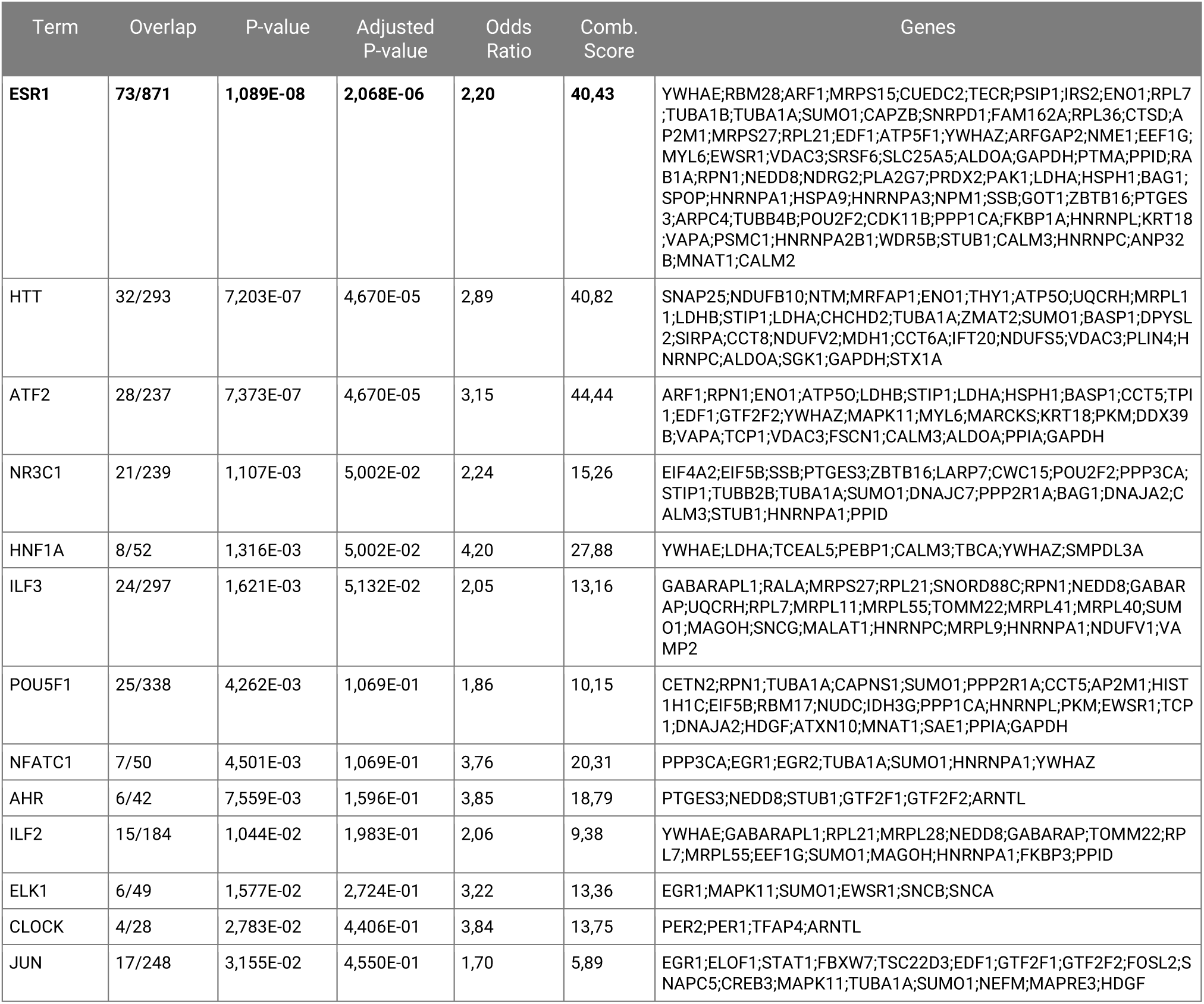
Transcription Factor Protein-Protein Interaction (TF PPI) analysis for the amygdala DEGs for <2 criteria vs alcohol-naive mice.

**Supplementary Table 3.**
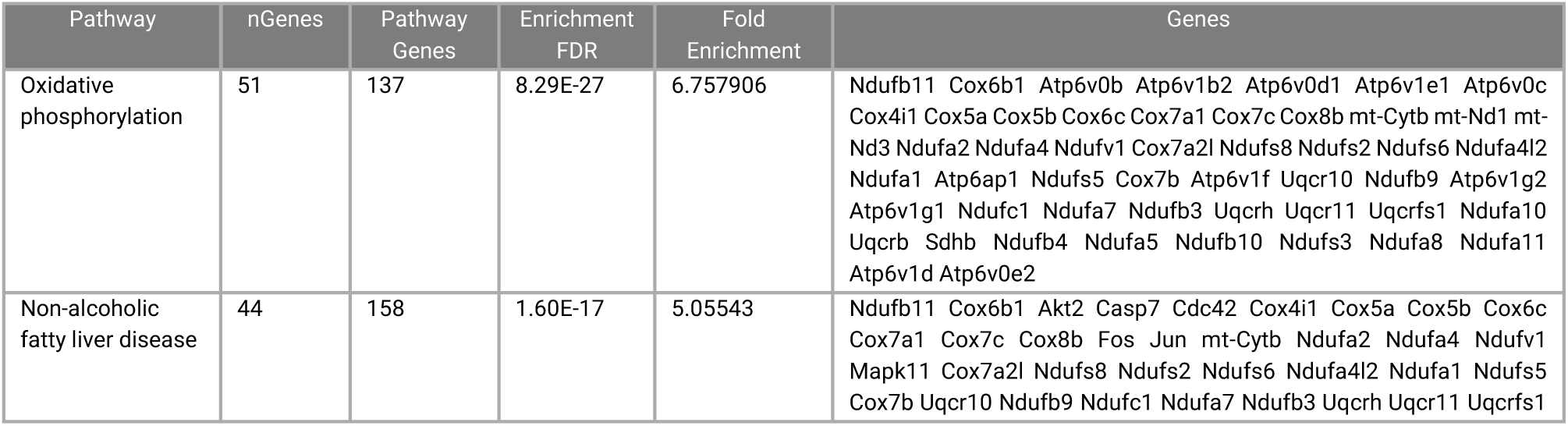
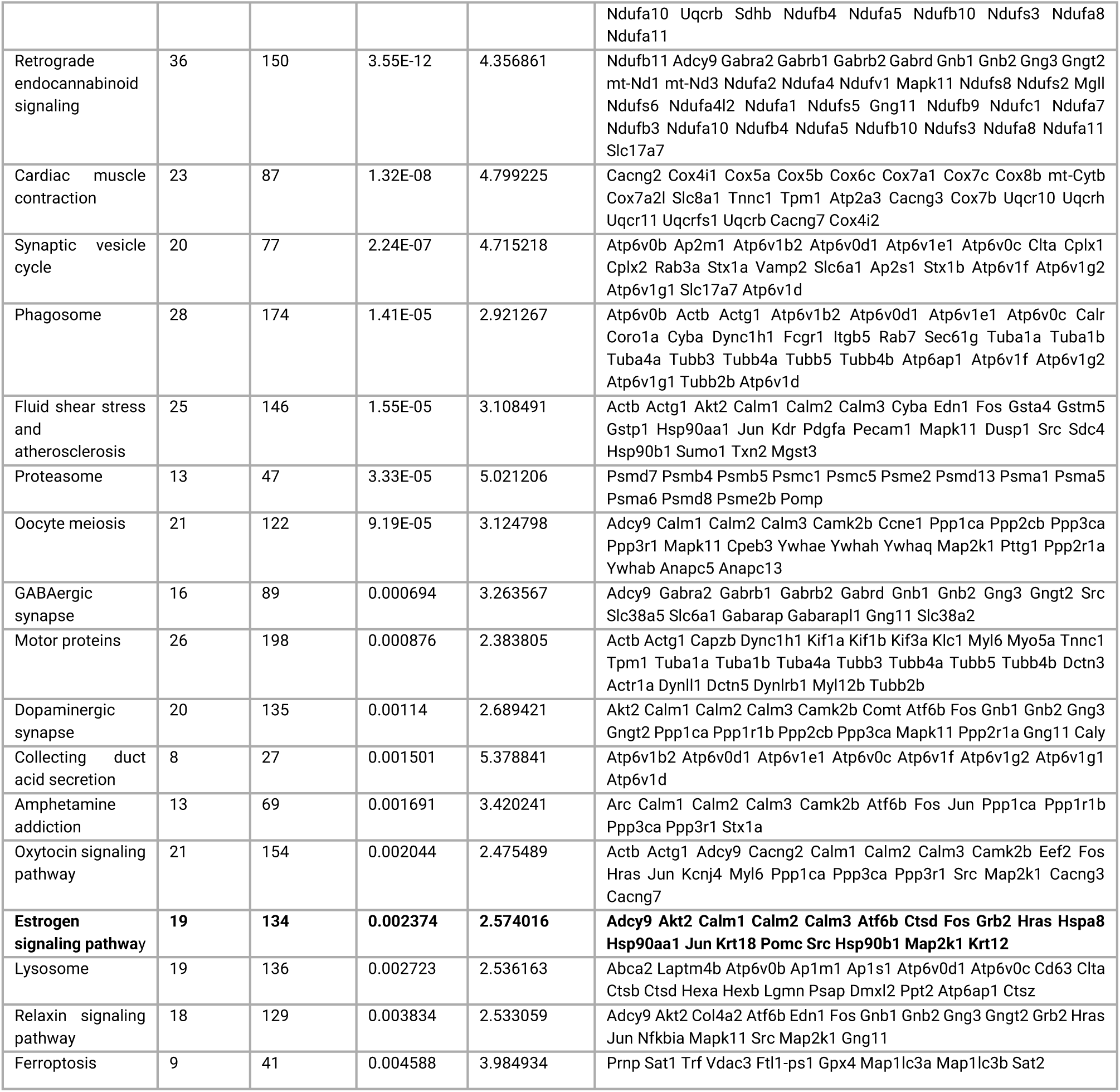
KEGG pathway enrichment analysis for the amygdala DEGs for ≥2 criteria vs alcohol-naive mice.

**Supplementary Table 4.**
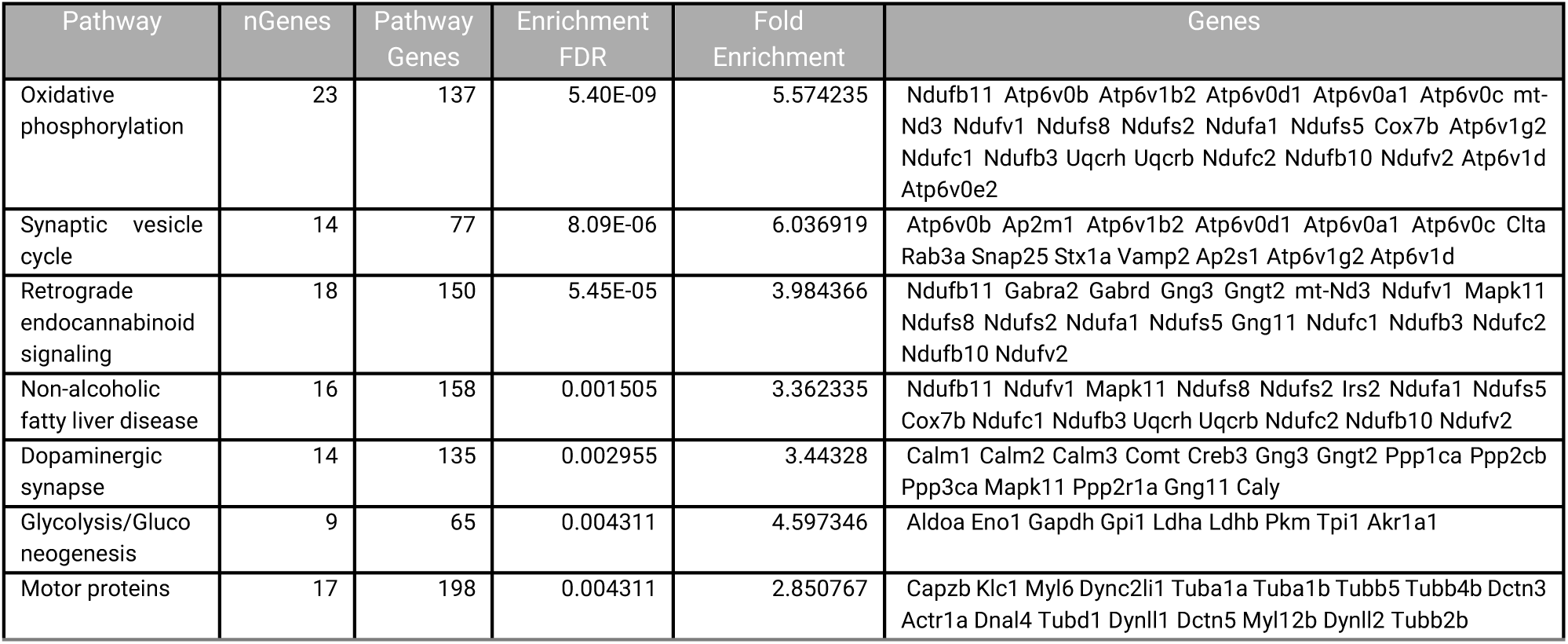
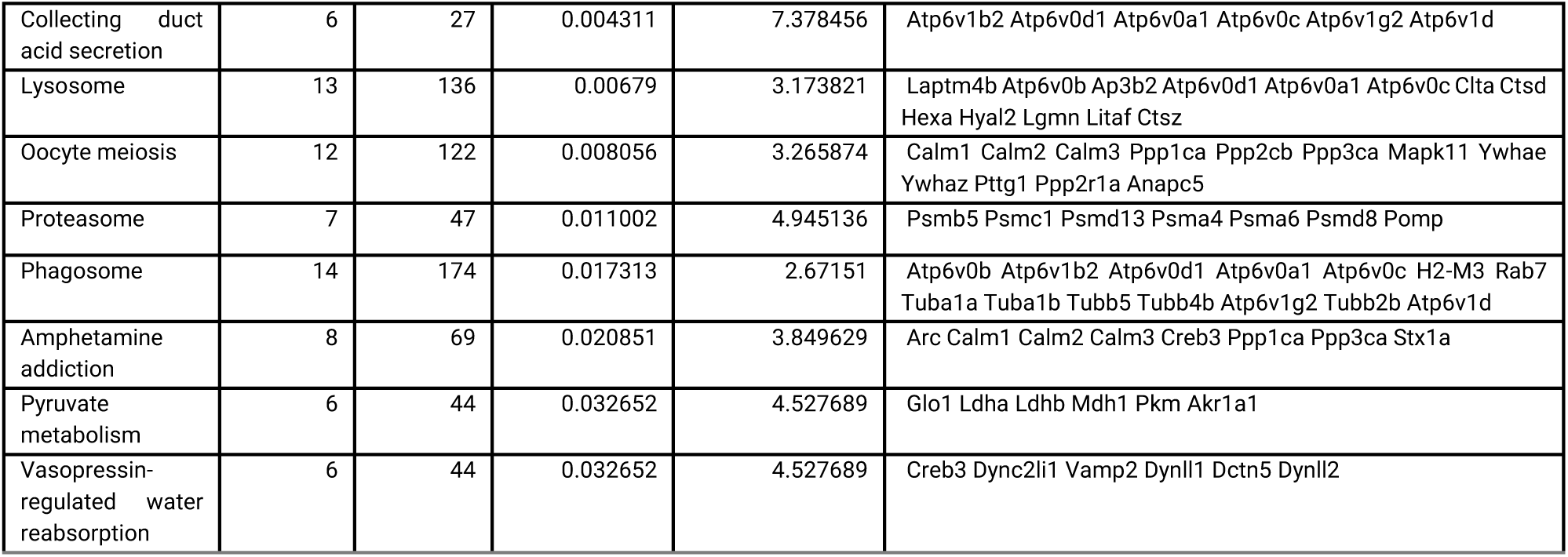
KEGG pathway enrichment analysis for the amygdala DEGs for <2 criteria vs alcohol-naive mice.

**Supplementary Table 5.** Associations of the rs6902771 *ESR1* genotypes with phenotypes of alcohol use disorder (AUD)

| | $\beta$ | 95% CI | OR | F | df1, df2 | p |
| --- | --- | --- | --- | --- | --- | --- |
| **Binge drinking days** |  |  |  |  |  |  |
| N[cases] = 200 |  |  |  |  |  |  |
| N[observations] = 40,333 |  |  |  |  |  |  |
| rs6902771=0 = 11,242 |  |  |  |  |  |  |
| rs6902771=1 = 20,111 |  |  |  |  |  |  |
| rs6902771=2 = 8,980 |  |  |  |  |  |  |
| AUD criteria | 0.0583 | [-0.0636, 0.1801] | 1.0600 | 0.891 | (1, 174) | 0.347 |
| Weekend days versus weekdays |  |  |  | 662.753 | (1, 40,322) | < 0.001 |
| Weekend days | 0.6972 | [0.6441, 0.7503] | 2.0081 |  |  |  |
| Weekdays | [Reference] |  |  |  |  |  |
| No lockdown versus lockdown |  |  |  | 6.950 | (1, 40,322) | 0.008 |
| No lockdown | 0.1484 | [0.0381, 0.2588] | 1.1600 |  |  |  |
| Lockdown | [Reference] |  |  |  |  |  |
| rs6902771 | 0.0122 | [-0.2745, 0.2989] | 1.0123 | 0.007 | (1, 180) | 0.933 |
| Day | -0.0015 | [-0.0020, -0.0010] | 0.9985 | 37.655 | (1, 40,322) | < 0.001 |
| rs6902771-by-day | 0.0008 | [0.0004, 0.0012] | 1.0008 | 18.355 | (1, 40,322) | < 0.001 |
| PC1 | -2.9252 | [-7.6756, 1.8252] | 0.0537 | 1.478 | (1, 169) | 0.226 |
| PC2 | 1.7332 | [-3.2652, 6.7317] | 5.6588 | 0.469 | (1, 168) | 0.495 |
| PC3 | -2.1915 | [-7.3587, 2.9758] | 0.1118 | 0.701 | (1, 168) | 0.404 |
| PC4 | -3.4346 | [-8.3730, 1.5038] | 0.0322 | 1.885 | (1, 171) | 0.172 |
| **Days with any alcohol use** |  |  |  |  |  |  |
| N[cases] = 200 |  |  |  |  |  |  |
| N[observations] = 40,333 |  |  |  |  |  |  |
| rs6902771=0 = 11,242 |  |  |  |  |  |  |
| rs6902771=1 = 20,111 |  |  |  |  |  |  |
| rs6902771=2 = 8,980 |  |  |  |  |  |  |
| AUD criteria | -0.0024 | [-0.1408, 0.1360] | 0.9976 | 0.001 | (1, 170) | 0.972 |
| Weekend days versus weekdays |  |  |  | 596.368 | (1, 40,322) | < 0.001 |
| Weekend days | 0.5805 | [0.5339, 0.6271] | 1.7869 |  |  |  |
| Weekdays | [Reference] |  |  |  |  |  |
| No lockdown versus lockdown |  |  |  | 48.728 | (1, 40,322) | < 0.001 |
| No lockdown | 0.3581 | [0.2576, 0.4587] | 1.4306 |  |  |  |
| Lockdown | [Reference] |  |  |  |  |  |
| rs6902771 | 0.0994 | [-0.2254, 0.4242] | 1.1045 | 0.365 | (1, 172) | 0.547 |
| Day | -0.0014 | [-0.0018, -0.0010] | 0.9986 | 46.965 | (1, 40,322) | < 0.001 |
| rs6902771-by-day | 0.0003 | [0.0000, 0.0007] | 1.0003 | 4.070 | (1, 40,322) | 0.044 |
| PC1 | -3.9723 | [-9.3847, 1.4402] | 0.0188 | 2.100 | (1, 165) | 0.149 |
| PC2 | 0.2557 | [-5.4326, 5.9441] | 1.2914 | 0.008 | (1, 163) | 0.929 |
| PC3 | -2.0495 | [-7.9385, 3.8395] | 0.1288 | 0.472 | (1, 164) | 0.493 |
| PC4 | 0.2209 | [-5.4101, 5.8520] | 1.2472 | 0.006 | (1, 168) | 0.938 |
| **Craving** |  |  |  |  |  |  |
| N[cases] = 195 |  |  |  |  |  |  |
| N[observations] = 16,364 |  |  |  |  |  |  |
| rs6902771=0 = 4,364 |  |  |  |  |  |  |
| rs6902771=1 = 8,346 |  |  |  |  |  |  |
| rs6902771=2 = 3,654 |  |  |  |  |  |  |
| AUD criteria | 0.1068 | [0.0242, 0.1894] |  | 6.503 | (1, 201) | 0.012 |
| Weekend days versus weekdays |  |  |  | 40.081 | (1, 16,189) | < 0.001 |
| Weekend days | 0.1004 | [0.0693, 0.1315] |  |  |  |  |
| Weekdays | [Reference] |  |  |  |  |  |
| No lockdown versus lockdown |  |  |  | 1.185 | (1, 16,275) | 0.276 |
| No lockdown | 0.0354 | [-0.0284, 0.0991] |  |  |  |  |
| Lockdown | [Reference] |  |  |  |  |  |
| rs6902771 | 0.1083 | [-0.0850, 0.3015] |  | 1.220 | (1, 208) | 0.271 |
| Day | -0.0009 | [-0.0012, -0.0007] |  | 41.527 | (1, 16,287) | < 0.001 |
| rs6902771-by-day | 0.0005 | [0.0003, 0.0008] |  | 22.002 | (1, 16,275) | < 0.001 |
| PC1 | -0.8838 | [-4.0829, 2.3153] |  | 0.297 | (1, 197) | 0.586 |
| PC2 | -1.3640 | [-4.7325, 2.0045] |  | 0.638 | (1, 196) | 0.426 |
| PC3 | -2.4378 | [-5.9268, 1.0511] |  | 1.899 | (1, 196) | 0.170 |
| PC4 | -0.6677 | [-4.0120, 2.6765] |  | 0.155 | (1, 199) | 0.694 |
| **Loss of control**<br>N[cases] = 184<br>N[observations] = 3,975<br>rs6902771=0 = 1,059<br>rs6902771=1 = 2,046<br>rs6902771=2 = 870 |  |  |  |  |  |  |
| AUD criteria | 0.1424 | [0.0685, 0.2163] |  | 14.428 | (1, 193) | < 0.001 |
| Weekend days versus weekdays |  |  |  | 0.010 | (1, 3,819) | 0.919 |
| Weekend days | 0.0027 | [-0.0496, 0.0550] |  |  |  |  |
| Weekdays | [Reference] |  |  |  |  |  |
| No lockdown versus lockdown |  |  |  | 0.442 | (1, 3,890) | 0.506 |
| No lockdown | 0.0358 | [-0.0699, 0.1416] |  |  |  |  |
| Lockdown | [Reference] |  |  |  |  |  |
| rs6902771 | -0.0259 | [-0.2035, 0.1517] |  | 0.083 | (1, 230) | 0.774 |
| Day | -0.0010 | [-0.0015, -0.0005] |  | 17.389 | (1, 3,890) | < 0.001 |
| rs6902771-by-day | 0.0006 | [0.0002, 0.0010] |  | 9.104 | (1, 3,874) | 0.003 |
| PC1 | -2.9514 | [-5.7612, -0.1415] |  | 4.292 | (1, 191) | 0.040 |
| PC2 | -0.9837 | [-3.9549, 1.9876] |  | 0.427 | (1, 186) | 0.514 |
| PC3 | -0.6176 | [-3.6755, 2.4402] |  | 0.159 | (1, 183) | 0.691 |
| PC4 | -1.1030 | [-4.0558, 1.8498] |  | 0.543 | (1, 188) | 0.462 |
| The models include fixed and random intercepts. AUD, alcohol use disorder; 95% CI, 95% confidence interval; $\beta$, regression coefficient; OR, odds ratio; PC principal component. rs6902771 is coded as "0" for CC, "1" for CT, and "2" for TT. | | | | | | |

**Supplementary Table 6.** Associations of the rs11155819 *ESR1* genotypes with phenotypes of alcohol use disorder (AUD)

| | $\beta$ | 95% CI | OR | F | df1, df2 | p |
| --- | --- | --- | --- | --- | --- | --- |
| **Binge drinking days**<br>N[cases] = 198<br>N[observations] = 40,189<br>rs11155819=0 = 17,312<br>rs11155819=1 = 18,889<br>rs11155819=2 = 3,988 |  |  |  |  |  |  |
| AUD criteria | 0.0619 | [-0.0621, 0.1858] | 1.0638 | 0.971 | (1, 172) | 0.326 |
| Weekend days versus weekdays |  |  |  | 659.274 | (1, 40,178) | < 0.001 |
| Weekend days | 0.6969 | [0.6437, 0.7501] | 2.0075 |  |  |  |
| Weekdays | [Reference] |  |  |  |  |  |
| No lockdown versus lockdown |  |  |  | 6.534 | (1, 40,178) | 0.011 |
| No lockdown | 0.1444 | [0.0337, 0.2550] | 1.1553 |  |  |  |
| Lockdown | [Reference] |  |  |  |  |  |
| rs11155819 | -0.0495 | [-0.3724, 0.2734] | 0.9517 | 0.091 | (1, 183) | 0.763 |
| Day | -0.0013 | [-0.0017, -0.0008] | 0.9987 | 36.298 | (1, 40,178) | < 0.001 |
| rs11155819-by-day | 0.0008 | [0.0004, 0.0012] | 1.0008 | 15.981 | (1, 40,178) | < 0.001 |
| PC1 | -2.5699 | [-7.4597, 2.3199] | 0.0765 | 1.077 | (1, 166) | 0.301 |
| PC2 | 1.7185 | [-3.3340, 6.7710] | 5.5763 | 0.451 | (1, 167) | 0.503 |
| PC3 | -2.2971 | [-7.4974, 2.9033] | 0.1006 | 0.761 | (1, 166) | 0.384 |
| PC4 | -3.3506 | [-8.3067, 1.6055] | 0.0351 | 1.781 | (1, 169) | 0.184 |
| **Days with any alcohol use**<br>N[cases] = 198<br>N[observations] = 40,189<br>rs11155819=0 = 17,312<br>rs11155819=1 = 18,889<br>rs11155819=2 = 3,988 |  |  |  |  |  |  |
| AUD criteria | 0.0166 | [-0.1239, 0.1572] | 1.0168 | 0.055 | (1, 169) | 0.815 |
| Weekend days versus weekdays |  |  |  | 597.780 | (1, 40,178) | < 0.001 |
| Weekend days | 0.5821 | [0.5354, 0.6288] | 1.7898 |  |  |  |
| Weekdays | [Reference] |  |  |  |  |  |
| No lockdown versus lockdown |  |  |  | 45.196 | (1, 40,178) | < 0.001 |
| No lockdown | 0.3460 | [0.2451, 0.4469] | 1.4134 |  |  |  |
| Lockdown | [Reference] |  |  |  |  |  |
| rs11155819 | -0.2262 | [-0.5897, 0.1373] | 0.7976 | 1.508 | (1, 175) | 0.221 |
| Day | -0.0018 | [-0.0022, -0.0015] | 0.9982 | 107.506 | (1, 40,178) | < 0.001 |
| rs11155819-by-day | 0.0011 | [0.0008, 0.0015] | 1.0011 | 36.681 | (1, 40,178) | < 0.001 |
| PC1 | -3.9988 | [-9.5660, 1.5684] | 0.0183 | 2.012 | (1, 163) | 0.158 |
| PC2 | -0.0342 | [-5.7669, 5.6985] | 0.9664 | 0.000 | (1, 161) | 0.991 |
| PC3 | -2.3639 | [-8.2701, 3.5424] | 0.0941 | 0.625 | (1, 162) | 0.430 |
| PC4 | 0.3085 | [-5.3258, 5.9428] | 1.3614 | 0.012 | (1, 166) | 0.914 |
| **Craving**<br>N[cases] = 193<br>N[observations] = 16,327<br>rs11155819=0 = 7,210<br>rs11155819=1 = 7,432<br>rs11155819=2 = 1,685 |  |  |  |  |  |  |
| AUD criteria | 0.1075 | [0.0243, 0.1908] |  | 6.487 | (1, 198) | 0.012 |
| Weekend days versus weekdays |  |  |  | 39.848 | (1, 16,154) | < 0.001 |
| Weekend days | 0.1003 | [0.0691, 0.1314] |  |  |  |  |
| Weekdays | [Reference] |  |  |  |  |  |
| No lockdown versus lockdown |  |  |  | 1.013 | (1, 16,239) | 0.314 |
| No lockdown | 0.0328 | [-0.0310, 0.0966] |  |  |  |  |
| Lockdown | [Reference] |  |  |  |  |  |
| rs11155819 | 0.1190 | [-0.0965, 0.3345] |  | 1.185 | (1, 210) | 0.278 |
| Day | -0.0007 | [-0.0010, -0.0005] |  | 38.270 | (1, 16,226) | < 0.001 |
| rs11155819-by-day | 0.0005 | [0.0003, 0.0008] |  | 17.207 | (1, 16,245) | < 0.001 |
| PC1 | -0.4736 | [-3.7334, 2.7862] |  | 0.082 | (1, 194) | 0.775 |
| PC2 | -1.1466 | [-4.5166, 2.2233] |  | 0.450 | (1, 195) | 0.503 |
| PC3 | -2.4195 | [-5.8913, 1.0523] |  | 1.889 | (1, 194) | 0.171 |
| PC4 | -0.5651 | [-3.8815, 2.7513] |  | 0.113 | (1, 197) | 0.737 |
| **Loss of control**<br>N[cases] = 182<br>N[observations] = 3,965<br>rs11155819=0 = 1,778<br>rs11155819=1 = 1,794<br>rs11155819=2 = 393 |  |  |  |  |  |  |
| AUD criteria | 0.1443 | [0.0695, 0.2190] |  | 14.477 | (1, 190) | < 0.001 |
| Weekend days versus weekdays |  |  |  | 0.038 | (1, 3,811) | 0.846 |
| Weekend days | 0.0052 | [-0.0470, 0.0573] |  |  |  |  |
| Weekdays | [Reference] |  |  |  |  |  |
| No lockdown versus lockdown |  |  |  | 0.281 | (1, 3,880) | 0.596 |
| No lockdown | 0.0285 | [-0.0769, 0.1339] |  |  |  |  |
| Lockdown | [Reference] |  |  |  |  |  |
| rs11155819 | -0.1075 | [-0.3079, 0.0929] |  | 1.117 | (1, 222) | 0.292 |
| Day | -0.0012 | [-0.0016, -0.0008] |  | 36.183 | (1, 3,856) | < 0.001 |
| rs11155819-by-day | 0.0012 | [0.0008, 0.0016] |  | 31.362 | (1, 3,860) | < 0.001 |
| PC1 | -3.2161 | [-6.0886, -0.3435] |  | 4.878 | (1, 187) | 0.028 |
| PC2 | -0.7488 | [-3.7419, 2.2442] |  | 0.244 | (1, 185) | 0.622 |
| PC3 | -0.5387 | [-3.6032, 2.5258] |  | 0.120 | (1, 182) | 0.729 |
| PC4 | -0.9503 | [-3.8936, 1.9930] |  | 0.406 | (1, 187) | 0.525 |
| The models include fixed and random intercepts. AUD, alcohol use disorder; 95% CI, 95% confidence interval; $\beta$, regression coefficient; OR, odds ratio; PC principal component. rs11155819 is coded as "0" for TT, "1" for TC, and "2" for CC. | | | | | | |

**Supplementary Table 7.** Associations of the rs6557171 *ESR1* genotypes with phenotypes of alcohol use disorder (AUD)

| | $\beta$ | 95% CI | OR | F | df1, df2 | p |
| --- | --- | --- | --- | --- | --- | --- |
| **Binge drinking days**<br>N[cases] = 200<br>N[observations] = 40,333<br>rs6557171=0 = 19,908<br>rs6557171=1 = 16,901<br>rs6557171=2 = 3,524 |  |  |  |  |  |  |
| AUD criteria | 0.0655 | [-0.0564, 0.1873] | 1.0676 | 1.124 | (1, 175) | 0.291 |
| Weekend days versus weekdays |  |  |  | 663.565 | (1, 40,322) | < 0.001 |
| Weekend days | 0.6977 | [0.6446, 0.7507] | 2.0090 |  |  |  |
| Weekdays | [Reference] |  |  |  |  |  |
| No lockdown versus lockdown |  |  |  | 7.857 | (1, 40,322) | 0.005 |
| No lockdown | 0.1586 | [0.0477, 0.2695] | 1.1719 |  |  |  |
| Lockdown | [Reference] |  |  |  |  |  |
| rs6557171 | 0.1538 | [-0.1622, 0.4697] | 1.1662 | 0.922 | (1, 188) | 0.338 |
| Day | -0.0001 | [-0.0004, 0.0003] | 0.9999 | 0.203 | (1, 40,322) | 0.652 |
| rs6557171-by-day | -0.0010 | [-0.0014, -0.0005] | 0.9990 | 20.800 | (1, 40,322) | < 0.001 |
| PC1 | -2.9247 | [-7.6817, 1.8323] | 0.0537 | 1.473 | (1, 170) | 0.227 |
| PC2 | 1.7859 | [-3.2253, 6.7972] | 5.9651 | 0.495 | (1, 169) | 0.483 |
| PC3 | -2.3131 | [-7.5205, 2.8943] | 0.0990 | 0.769 | (1, 168) | 0.382 |
| PC4 | -3.1777 | [-8.1113, 1.7560] | 0.0417 | 1.616 | (1, 171) | 0.205 |
| **Days with any alcohol use**<br>N[cases] = 200<br>N[observations] = 40,333<br>rs6557171=0 = 19,908<br>rs6557171=1 = 16,901<br>rs6557171=2 = 3,524 |  |  |  |  |  |  |
| AUD criteria | 0.0054 | [-0.1332, 0.1440] | 1.0054 | 0.006 | (1, 171) | 0.939 |
| Weekend days versus weekdays |  |  |  | 597.146 | (1, 40,322) | < 0.001 |
| Weekend days | 0.5811 | [0.5345, 0.6277] | 1.7881 |  |  |  |
| Weekdays | [Reference] |  |  |  |  |  |
| No lockdown versus lockdown |  |  |  | 47.575 | (1, 40,322) | < 0.001 |
| No lockdown | 0.3541 | [0.2535, 0.4547] | 1.4249 |  |  |  |
| Lockdown | [Reference] |  |  |  |  |  |
| rs6557171 | 0.1590 | [-0.1988, 0.5168] | 1.1723 | 0.769 | (1, 181) | 0.382 |
| Day | -0.0004 | [-0.0008, -0.0001] | 0.9996 | 7.777 | (1, 40,322) | 0.005 |
| rs6557171-by-day | -0.0011 | [-0.0015, -0.0007] | 0.9989 | 35.299 | (1, 40,322) | < 0.001 |
| PC1 | -3.9839 | [-9.4080, 1.4403] | 0.0186 | 2.103 | (1, 165) | 0.149 |
| PC2 | 0.3494 | [-5.3567, 6.0556] | 1.4183 | 0.015 | (1, 163) | 0.904 |
| PC3 | -2.1789 | [-8.1178, 3.7601] | 0.1132 | 0.525 | (1, 164) | 0.470 |
| PC4 | 0.4678 | [-5.1639, 6.0995] | 1.5965 | 0.027 | (1, 168) | 0.870 |
| **Craving**<br>N[cases] = 195<br>N[observations] = 16,364<br>rs6557171=0 = 8,156<br>rs6557171=1 = 6,707<br>rs6557171=2 = 1,501 |  |  |  |  |  |  |
| AUD criteria | 0.1246 | [0.0417, 0.2074] |  | 8.792 | (1, 201) | 0.003 |
| Weekend days versus weekdays |  |  |  | 40.049 | (1, 16,189) | < 0.001 |
| Weekend days | 0.1005 | [0.0694, 0.1316] |  |  |  |  |
| Weekdays | [Reference] |  |  |  |  |  |
| No lockdown versus lockdown |  |  |  | 1.308 | (1, 16,275) | 0.253 |
| No lockdown | 0.0372 | [-0.0266, 0.1010] |  |  |  |  |
| Lockdown | [Reference] |  |  |  |  |  |
| rs6557171 | -0.1144 | [-0.3312, 0.1025] |  | 1.080 | (1, 212) | 0.300 |
| Day | -0.0003 | [-0.0005, -0.0001] |  | 6.365 | (1, 16,257) | 0.012 |
| rs6557171-by-day | -0.0002 | [-0.0004, 0.0001] |  | 1.865 | (1, 16,277) | 0.172 |
| PC1 | -0.9969 | [-4.2030, 2.2091] |  | 0.376 | (1, 197) | 0.540 |
| PC2 | -1.1697 | [-4.5496, 2.2102] |  | 0.466 | (1, 196) | 0.496 |
| PC3 | -2.2998 | [-5.8212, 1.2217] |  | 1.659 | (1, 197) | 0.199 |
| PC4 | -0.4078 | [-3.7464, 2.9307] |  | 0.058 | (1, 199) | 0.810 |
| **Loss of control**<br>N[cases] = 184<br>N[observations] = 3,975<br>rs6557171=0 = 2,002<br>rs6557171=1 = 1,612<br>rs6557171=2 = 361 |  |  |  |  |  |  |
| AUD criteria | 0.1410 | [0.0671, 0.2149] |  | 14.175 | (1, 193) | < 0.001 |
| Weekend days versus weekdays |  |  |  | 0.005 | (1, 3,819) | 0.941 |
| Weekend days | 0.0020 | [-0.0504, 0.0543] |  |  |  |  |
| Weekdays | [Reference] |  |  |  |  |  |
| No lockdown versus lockdown |  |  |  | 0.504 | (1, 3,891) | 0.478 |
| No lockdown | 0.0383 | [-0.0675, 0.1441] |  |  |  |  |
| Lockdown | [Reference] |  |  |  |  |  |
| rs6557171 | 0.1479 | [-0.0527, 0.3485] |  | 2.111 | (1, 229) | 0.148 |
| Day | -0.0002 | [-0.0006, 0.0001] |  | 1.513 | (1, 3,851) | 0.219 |
| rs6557171-by-day | -0.0003 | [-0.0007, 0.0001] |  | 2.433 | (1, 3,867) | 0.119 |
| PC1 | -2.9392 | [-5.7422, -0.1362] |  | 4.278 | (1, 191) | 0.040 |
| PC2 | -1.0402 | [-4.0067, 1.9263] |  | 0.479 | (1, 186) | 0.490 |
| PC3 | -0.8841 | [-3.9597, 2.1915] |  | 0.322 | (1, 184) | 0.571 |
| PC4 | -1.0103 | [-3.9403, 1.9197] |  | 0.463 | (1, 188) | 0.497 |
| The models include fixed and random intercepts. AUD, alcohol use disorder; 95% CI, 95% confidence interval; $\beta$, regression coefficient; OR, odds ratio; PC principal component. rs6557171 is coded as "0" for CC, "1" for CT, and "2" for TT. | | | | | | |

**Supplementary Table 8.** Associations of the rs2982712 *ESR1* genotypes with phenotypes of alcohol use disorder (AUD)

| | $\beta$ | 95% CI | OR | F | df1, df2 | p |
| --- | --- | --- | --- | --- | --- | --- |
| **Binge drinking days** |  |  |  |  |  |  |
| N[cases] = 200 |  |  |  |  |  |  |
| N[observations] = 40,333 |  |  |  |  |  |  |
| rs2982712=0 = 12,429 |  |  |  |  |  |  |
| rs2982712=1 = 18,259 |  |  |  |  |  |  |
| rs2982712=2 = 9,645 |  |  |  |  |  |  |
| AUD criteria | 0.0625 | [-0.0587, 0.1836] | 1.0644 | 1.036 | (1, 174) | 0.310 |
| Weekend days versus weekdays |  |  |  | 663.507 | (1, 40,322) | < 0.001 |
| Weekend days | 0.6974 | [0.6443, 0.7505] | 2.0085 |  |  |  |
| Weekdays | [Reference] |  |  |  |  |  |
| No lockdown versus lockdown |  |  |  | 7.283 | (1, 40,322) | 0.007 |
| No lockdown | 0.1522 | [0.0417, 0.2628] | 1.1644 |  |  |  |
| Lockdown | [Reference] |  |  |  |  |  |
| rs2982712 | -0.1038 | [-0.3848, 0.1772] | 0.9014 | 0.531 | (1, 186) | 0.467 |
| Day | -0.0005 | [-0.0009, -0.0001] | 0.9995 | 5.429 | (1, 40,322) | 0.020 |
| rs2982712-by-day | -0.0002 | [-0.0005, 0.0002] | 0.9998 | 0.714 | (1, 40,322) | 0.398 |
| PC1 | -2.8366 | [-7.5934, 1.9201] | 0.0586 | 1.386 | (1, 169) | 0.241 |
| PC2 | 1.5070 | [-3.5231, 6.5370] | 4.5130 | 0.350 | (1, 168) | 0.555 |
| PC3 | -2.0023 | [-7.1898, 3.1853] | 0.1350 | 0.581 | (1, 168) | 0.447 |
| PC4 | -3.1561 | [-8.0758, 1.7635] | 0.0426 | 1.604 | (1, 171) | 0.207 |
| **Days with any alcohol use** |  |  |  |  |  |  |
| N[cases] = 200 |  |  |  |  |  |  |
| N[observations] = 40,333 |  |  |  |  |  |  |
| rs2982712=0 = 12,429 |  |  |  |  |  |  |
| rs2982712=1 = 18,259 |  |  |  |  |  |  |
| rs2982712=2 = 9,645 |  |  |  |  |  |  |
| AUD criteria | 0.0064 | [-0.1315, 0.1443] | 1.0064 | 0.008 | (1, 171) | 0.927 |
| Weekend days versus weekdays |  |  |  | 596.700 | (1, 40,322) | < 0.001 |
| Weekend days | 0.5806 | [0.5340, 0.6272] | 1.7871 |  |  |  |
| Weekdays | [Reference] |  |  |  |  |  |
| No lockdown versus lockdown |  |  |  | 49.072 | (1, 40,322) | < 0.001 |
| No lockdown | 0.3594 | [0.2589, 0.4600] | 1.4325 |  |  |  |
| Lockdown | [Reference] |  |  |  |  |  |
| rs2982712 | -0.0131 | [-0.3308, 0.3047] | 0.9870 | 0.007 | (1, 176) | 0.935 |
| Day | -0.0012 | [-0.0016, -0.0008] | 0.9988 | 35.058 | (1, 40,322) | < 0.001 |
| rs2982712-by-day | 0.0001 | [-0.0002, 0.0004] | 1.0001 | 0.459 | (1, 40,322) | 0.498 |
| PC1 | -4.0474 | [-9.4798, 1.3851] | 0.0175 | 2.164 | (1, 165) | 0.143 |
| PC2 | 0.3120 | [-5.4266, 6.0506] | 1.3662 | 0.012 | (1, 163) | 0.915 |
| PC3 | -2.2052 | [-8.1291, 3.7187] | 0.1102 | 0.540 | (1, 165) | 0.463 |
| PC4 | 0.4698 | [-5.1519, 6.0915] | 1.5997 | 0.027 | (1, 168) | 0.869 |
| **Craving** |  |  |  |  |  |  |
| N[cases] = 195 |  |  |  |  |  |  |
| N[observations] = 16,364 |  |  |  |  |  |  |
| rs2982712=0 = 5,100 |  |  |  |  |  |  |
| rs2982712=1 = 7,380 |  |  |  |  |  |  |
| rs2982712=2 = 3,884 |  |  |  |  |  |  |
| AUD criteria | 0.1189 | [0.0361, 0.2016] |  | 8.026 | (1, 200) | 0.005 |
| Weekend days versus weekdays |  |  |  | 40.061 | (1, 16,189) | < 0.001 |
| Weekend days | 0.1005 | [0.0694, 0.1316] |  |  |  |  |
| Weekdays | [Reference] |  |  |  |  |  |
| No lockdown versus lockdown |  |  |  | 1.303 | (1, 16,274) | 0.254 |
| No lockdown | 0.0372 | [-0.0266, 0.1010] |  |  |  |  |
| Lockdown | [Reference] |  |  |  |  |  |
| rs2982712 | 0.0301 | [-0.1606, 0.2209] |  | 0.097 | (1, 209) | 0.756 |
| Day | -0.0004 | [-0.0006, -0.0001] |  | 7.853 | (1, 16,272) | 0.005 |
| rs2982712-by-day | -0.0000 | [-0.0002, 0.0002] |  | 0.015 | (1, 16,261) | 0.902 |
| PC1 | -0.9957 | [-4.2204, 2.2291] |  | 0.371 | (1, 197) | 0.543 |
| PC2 | -1.2246 | [-4.6360, 2.1869] |  | 0.501 | (1, 196) | 0.480 |
| PC3 | -2.6529 | [-6.1829, 0.8772] |  | 2.196 | (1, 197) | 0.140 |
| PC4 | -0.3499 | [-3.7011, 3.0012] |  | 0.042 | (1, 199) | 0.837 |
| **Loss of control**<br>N[cases] = 184<br>N[observations] = 3,975<br>rs2982712=0 = 1,249<br>rs2982712=1 = 1,782<br>rs2982712=2 = 944 |  |  |  |  |  |  |
| AUD criteria | 0.1446 | [0.0713, 0.2180] |  | 15.137 | (1, 193) | < 0.001 |
| Weekend days versus weekdays |  |  |  | 0.005 | (1, 3,819) | 0.942 |
| Weekend days | 0.0019 | [-0.0504, 0.0543] |  |  |  |  |
| Weekdays | [Reference] |  |  |  |  |  |
| No lockdown versus lockdown |  |  |  | 0.481 | (1, 3,890) | 0.488 |
| No lockdown | 0.0374 | [-0.0684, 0.1433] |  |  |  |  |
| Lockdown | [Reference] |  |  |  |  |  |
| rs2982712 | -0.0430 | [-0.2177, 0.1316] |  | 0.236 | (1, 231) | 0.628 |
| Day | -0.0002 | [-0.0007, 0.0002] |  | 1.259 | (1, 3,866) | 0.262 |
| rs2982712-by-day | -0.0002 | [-0.0005, 0.0002] |  | 1.046 | (1, 3,863) | 0.307 |
| PC1 | -2.9001 | [-5.7112, -0.0890] |  | 4.141 | (1, 191) | 0.043 |
| PC2 | -1.1100 | [-4.0933, 1.8734] |  | 0.539 | (1, 186) | 0.464 |
| PC3 | -0.4806 | [-3.5582, 2.5969] |  | 0.095 | (1, 184) | 0.758 |
| PC4 | -0.9861 | [-3.9189, 1.9468] |  | 0.440 | (1, 188) | 0.508 |
| The models include fixed and random intercepts. AUD, alcohol use disorder; 95% CI, 95% confidence interval; $\beta$, regression coefficient; OR, odds ratio; PC principal component. rs2982712 is coded as "0" for TT, "1" for TC, and "2" for CC. | | | | | | |

**Supplementary Table 10.** Correlations between blood *ESR1* mRNA levels and demographic data in human AUD female patients (NOAH study).

|  | r | p |
| --- | --- | --- |
| **Control** |  |  |
| Age | -0.197 | 0.067 |
| BMI | -0.065 | 0.552 |
| AUDIT score | 0.000 | 0.998 |
| **Early abstinent patients** |  |  |
| Age | 0.024 | 0.831 |
| BMI | 0.028 | 0.818 |
| PACS score | 0.148 | 0.206 |

## LITERATURE

1. Grant BF, Goldstein RB, Saha TD, Chou SP, Jung J, Zhang H, et al. Epidemiology of *DSM-5* Alcohol Use Disorder: Results From the National Epidemiologic Survey on Alcohol and Related Conditions III. JAMA Psychiatry. 2015;72:757.

2. Carvalho AF, Heilig M, Perez A, Probst C, Rehm J. Alcohol use disorders. The Lancet. 2019;394:781–792.

3. DSM-5. https://www.psychiatry.org/psychiatrists/practice/dsm. Accessed 7 March 2022.

4. Lenz B, Derntl B. Sex-sensitive and gender-sensitive care for patients with mental disorders. Lancet Psychiatry. 2024:S2215–0366(24)00330-4.

5. White AM. Gender Differences in the Epidemiology of Alcohol Use and Related Harms in the United States. Alcohol Res. 2020;40:01.

6. Piazza NJ, Vrbka JL, Yeager RD. Telescoping of Alcoholism in Women Alcoholics. International Journal of the Addictions. 1989;24:19–28.

7. Evans SM, Reynolds B. Sex differences in drug abuse: Etiology, prevention, and treatment. Exp Clin Psychopharmacol. 2015;23:195–196.

8. Lynch WJ, Roth ME, Carroll ME. Biological basis of sex differences in drug abuse: preclinical and clinical studies. Psychopharmacology. 2002;164:121–137.

9. Mühle C, Barry B, Weinland C, Kornhuber J, Lenz B. Estrogen receptor 1 gene variants and estradiol activities in alcohol dependence. Prog Neuropsychopharmacol Biol Psychiatry. 2019;92:301–307.

10. Hoffmann S, Gerhardt S, Mühle C, Reinhard I, Reichert D, Bach P, et al. Associations of Menstrual Cycle and Progesterone-to-Estradiol Ratio With Alcohol Consumption in Alcohol Use Disorder: A Sex-Separated Multicenter Longitudinal Study. AJP. 2024;181:445–456.

11. Priddy BM, Carmack SA, Thomas LC, Vendruscolo JCM, Koob GF, Vendruscolo LF. Sex, strain, and estrous cycle influences on alcohol drinking in rats. Pharmacol Biochem Behav. 2017;152:61–67.

12. Roberts AJ, Smith AD, Weiss F, Rivier C, Koob GF. Estrous Cycle Effects on Operant Responding for Ethanol in Female Rats. Alcoholism: Clinical and Experimental Research. 1998;22:1564–1569.

13. Torres OV, Walker EM, Beas BS, O’Dell LE. Female Rats Display Enhanced Rewarding Effects of Ethanol That Are Hormone Dependent. Alcoholism: Clinical and Experimental Research. 2014;38:108–115.

14. Ford MM, Eldridge JC, Samson HH. Ethanol consumption in the female Long-Evans rat: a modulatory role of estradiol. Alcohol. 2002;26:103–113.

15. Handa RJ, Weiser MJ. Gonadal steroid hormones and the hypothalamo–pituitary–adrenal axis. Frontiers in Neuroendocrinology. 2014;35:197–220.

16. Mitra SW, Hoskin E, Yudkovitz J, Pear L, Wilkinson HA, Hayashi S, et al. Immunolocalization of Estrogen Receptor β in the Mouse Brain: Comparison with Estrogen Receptor α. Endocrinology. 2003;144:2055–2067.

17. Mitterling KL, Spencer JL, Dziedzic N, Shenoy S, McCarthy K, Waters EM, et al. Cellular and subcellular localization of estrogen and progestin receptor immunoreactivities in the mouse hippocampus. J Comp Neurol. 2010;518:2729–2743.

18. Osterlund M, Kuiper GG, Gustafsson JA, Hurd YL. Differential distribution and regulation of estrogen receptor-alpha and -beta mRNA within the female rat brain. Brain Res Mol Brain Res. 1998;54:175–180.

19. Le Moëne O, Stavarache M, Ogawa S, Musatov S, Ågmo A. Estrogen receptors α and β in the central amygdala and the ventromedial nucleus of the hypothalamus: Sociosexual behaviors, fear and arousal in female rats during emotionally challenging events. Behav Brain Res. 2019;367:128–142.

20. Miller CK, Krentzel AA, Meitzen J. ERα Stimulation Rapidly Modulates Excitatory Synapse Properties in Female Rat Nucleus Accumbens Core. Neuroendocrinology. 2023;113:1140–1153.

21. Smith CC, McMahon LL. Estrogen-induced increase in the magnitude of long-term potentiation occurs only when the ratio of NMDA transmission to AMPA transmission is increased. J Neurosci. 2005;25:7780–7791.

22. Woolley CS, McEwen BS. Estradiol regulates hippocampal dendritic spine density via an N-methyl-D-aspartate receptor-dependent mechanism. J Neurosci. 1994;14:7680–7687.

23. Gross KS, Mermelstein PG. Estrogen receptor signaling through metabotropic glutamate receptors. Vitam Horm. 2020;114:211–232.

24. Almey A, Milner TA, Brake WG. Estrogen receptors in the central nervous system and their implication for dopamine-dependent cognition in females. Horm Behav. 2015;74:125–138.

25. Lymer J, Bergman H, Yang S, Mallick R, Galea LAM, Choleris E, et al. The effects of estrogens on spatial learning and memory in female rodents – A systematic review and meta-analysis. Hormones and Behavior. 2024;164:105598.

26. Vandegrift BJ, Hilderbrand ER, Satta R, Tai R, He D, You C, et al. Estrogen Receptor α Regulates Ethanol Excitation of Ventral Tegmental Area Neurons and Binge Drinking in Female Mice. J Neurosci. 2020;40:5196–5207.

27. Zallar LJ, Rivera-Irizarry JK, Hamor PU, Pigulevskiy I, Rico Rozo A-S, Mehanna H, et al. Rapid nongenomic estrogen signaling controls alcohol drinking behavior in mice. Nat Commun. 2024;15:10725.

28. Chen H, Lu Y, Xiong R, Rosales CI, Coles C, Hamada K, et al. Effect of a brain-penetrant selective estrogen receptor degrader (SERD) on binge drinking in female mice. Alcoholism: Clinical and Experimental Research. 2022;46:1313–1320.

29. Treutlein J, Cichon S, Ridinger M, Wodarz N, Soyka M, Zill P, et al. Genome-wide Association Study of Alcohol Dependence. Arch Gen Psychiatry. 2009;66:773.

30. Hochgerner H, Singh S, Tibi M, Lin Z, Skarbianskis N, Admati I, et al. Neuronal types in the mouse amygdala and their transcriptional response to fear conditioning. Nat Neurosci. 2023;26:2237–2249.

31. Everitt BJ, Parkinson JA, Olmstead MC, Arroyo M, Robledo P, Robbins TW. Associative processes in addiction and reward. The role of amygdala-ventral striatal subsystems. Ann N Y Acad Sci. 1999;877:412–438.

32. Johnson ST, Grabenhorst F. The amygdala and the pursuit of future rewards. Front Neurosci. 2024;18:1517231.

33. Knapska E, Radwanska K, Werka T, Kaczmarek L. Functional internal complexity of amygdala: focus on gene activity mapping after behavioral training and drugs of abuse. Physiol Rev. 2007;87:1113–1173.

34. Stefaniuk M, Beroun A, Lebitko T, Markina O, Leski S, Meyza K, et al. Matrix Metalloproteinase-9 and Synaptic Plasticity in the Central Amygdala in Control of Alcohol-Seeking Behavior. Biological Psychiatry. 2017;81:907–917.

35. Pagano R, Salamian A, Zielinski J, Beroun A, Nalberczak-Skóra M, Skonieczna E, et al. Arc controls alcohol cue relapse by a central amygdala mechanism. Mol Psychiatry. 2023;28:733–745.

36. Radwanska K, Wrobel E, Korkosz A, Rogowski A, Kostowski W, Bienkowski P, et al. Alcohol Relapse Induced by Discrete Cues Activates Components of AP-1 Transcription Factor and ERK Pathway in the Rat Basolateral and Central Amygdala. Neuropsychopharmacology. 2008;33:1835–1846.

37. Grant S, London ED, Newlin DB, Villemagne VL, Liu X, Contoreggi C, et al. Activation of memory circuits during cue-elicited cocaine craving. PNAS. 1996;93:12040–12045.

38. Kruzich PJ, See RE. Differential Contributions of the Basolateral and Central Amygdala in the Acquisition and Expression of Conditioned Relapse to Cocaine-Seeking Behavior. J Neurosci. 2001;21:RC155.

39. Vafaie N, Kober H. Association of Drug Cues and Craving With Drug Use and Relapse: A Systematic Review and Meta-analysis. JAMA Psychiatry. 2022;79:641–650.

40. Radwanska K, Kaczmarek L. Characterization of an alcohol addiction-prone phenotype in mice: Characterization of an alcohol addiction-prone phenotype. Addiction Biology. 2012;17:601–612.

41. Beroun A, Nalberczak-Skóra M, Harda Z, Piechota M, Ziółkowska M, Cały A, et al. Generation of silent synapses in dentate gyrus correlates with development of alcohol addiction. Neuropsychopharmacol. 2018;43:1989–1999.

42. Salamian A, Pagano R, Skonieczna E, Kalinichenko LS, Puchalska M, Yiğit Ünlü G, et al. Hippocampal Cofilin and CFL1 gene variants are linked to Alcohol Use Disorder phenotypes. Mol Psychiatry. 2025. 11 September 2025. 10.1038/s41380-025-03226-3.

43. Ge SX, Jung D, Yao R. ShinyGO: a graphical gene-set enrichment tool for animals and plants. Bioinformatics. 2020;36:2628–2629.

44. Xie Z, Bailey A, Kuleshov MV, Clarke DJB, Evangelista JE, Jenkins SL, et al. Gene Set Knowledge Discovery with Enrichr. Curr Protoc. 2021;1:e90.

45. Chen EY, Tan CM, Kou Y, Duan Q, Wang Z, Meirelles GV, et al. Enrichr: interactive and collaborative HTML5 gene list enrichment analysis tool. BMC Bioinformatics. 2013;14:128.

46. Kuleshov MV, Jones MR, Rouillard AD, Fernandez NF, Duan Q, Wang Z, et al. Enrichr: a comprehensive gene set enrichment analysis web server 2016 update. Nucleic Acids Res. 2016;44:W90–97.

47. Heinz S, Benner C, Spann N, Bertolino E, Lin YC, Laslo P, et al. Simple combinations of lineage-determining transcription factors prime cis-regulatory elements required for macrophage and B cell identities. Mol Cell. 2010;38:576–589.

48. Subramanian A, Tamayo P, Mootha VK, Mukherjee S, Ebert BL, Gillette MA, et al. Gene set enrichment analysis: a knowledge-based approach for interpreting genome-wide expression profiles. Proc Natl Acad Sci U S A. 2005;102:15545–15550.

49. Liberzon A, Birger C, Thorvaldsdóttir H, Ghandi M, Mesirov JP, Tamayo P. The Molecular Signatures Database (MSigDB) hallmark gene set collection. Cell Syst. 2015;1:417–425.

50. Castanza AS, Recla JM, Eby D, Thorvaldsdóttir H, Bult CJ, Mesirov JP. Extending support for mouse data in the Molecular Signatures Database (MSigDB). Nat Methods. 2023;20:1619–1620.

51. Schindelin J, Arganda-Carreras I, Frise E, Kaynig V, Longair M, Pietzsch T, et al. Fiji: an open-source platform for biological-image analysis. Nat Methods. 2012;9:676–682.

52. McLean AC, Valenzuela N, Fai S, Bennett SAL. Performing vaginal lavage, crystal violet staining, and vaginal cytological evaluation for mouse estrous cycle staging identification. J Vis Exp. 2012:e4389.

53. Byers SL, Wiles MV, Dunn SL, Taft RA. Mouse Estrous Cycle Identification Tool and Images. PLOS ONE. 2012;7:e35538.

54. Paxinos and Franklin’s the Mouse Brain in Stereotaxic Coordinates, Compact - 5th Edition.

55. Hoffmann S, Gerhardt S, Mühle C, Reinhard I, Reichert D, Bach P, et al. Associations of Menstrual Cycle and Progesterone-to-Estradiol Ratio With Alcohol Consumption in Alcohol Use Disorder: A Sex-Separated Multicenter Longitudinal Study. Am J Psychiatry. 2024;181:445–456.

56. Spanagel R, Bach P, Banaschewski T, Beck A, Bermpohl F, Bernardi RE, et al. The ReCoDe addiction research consortium: Losing and regaining control over drug intake-Findings and future perspectives. Addict Biol. 2024;29:e13419.

57. Heinz A, Kiefer F, Smolka MN, Endrass T, Beste C, Beck A, et al. Addiction Research Consortium: Losing and regaining control over drug intake (ReCoDe)-From trajectories to mechanisms and interventions. Addict Biol. 2020;25:e12866.

58. Purcell S, Neale B, Todd-Brown K, Thomas L, Ferreira MAR, Bender D, et al. PLINK: a tool set for whole-genome association and population-based linkage analyses. Am J Hum Genet. 2007;81:559–575.

59. Loh P-R, Danecek P, Palamara PF, Fuchsberger C, A Reshef Y, K Finucane H, et al. Reference-based phasing using the Haplotype Reference Consortium panel. Nat Genet. 2016;48:1443–1448.

60. Das S, Forer L, Schönherr S, Sidore C, Locke AE, Kwong A, et al. Next-generation genotype imputation service and methods. Nat Genet. 2016;48:1284–1287.

61. Lenz B, Mühle C, Braun B, Weinland C, Bouna-Pyrrou P, Behrens J, et al. Prenatal and adult androgen activities in alcohol dependence. Acta Psychiatr Scand. 2017;136:96–107.

62. Weinland C, Mühle C, Kornhuber J, Lenz B. Body mass index and craving predict 24-month hospital readmissions of alcohol-dependent in-patients following withdrawal. Prog Neuropsychopharmacol Biol Psychiatry. 2019;90:300–307.

63. Brazdis R-M, von Zimmermann C, Lenz B, Kornhuber J, Mühle C. Peripheral Upregulation of Parkinson’s Disease-Associated Genes Encoding α-Synuclein, β-Glucocerebrosidase, and Ceramide Glucosyltransferase in Major Depression. Int J Mol Sci. 2024;25:3219.

64. Cavalcanti FN, Lucas TFG, Lazari MFM, Porto CS. Estrogen receptor ESR1 mediates activation of ERK1/2, CREB, and ELK1 in the corpus of the epididymis. J Mol Endocrinol. 2015;54:339–349.

65. Besnard A, Bouveyron N, Kappes V, Pascoli V, Pagès C, Heck N, et al. Alterations of Molecular and Behavioral Responses to Cocaine by Selective Inhibition of Elk-1 Phosphorylation. J Neurosci. 2011;31:14296–14307.

66. Kalet BT, Anglin SR, Handschy A, O’Donoghue LE, Halsey C, Chubb L, et al. Transcription factor Ets1 cooperates with estrogen receptor α to stimulate estradiol-dependent growth in breast cancer cells and tumors. PLoS One. 2013;8:e68815.

67. Gilpin NW, Herman MA, Roberto M. The central amygdala as an integrative hub for anxiety and alcohol use disorders. Biol Psychiatry. 2015;77:859–869.

68. Zhang X, Kim J, Tonegawa S. Amygdala Reward Neurons Form and Store Fear Extinction Memory. Neuron. 2020;105:1077–1093.e7.

69. Liu F, Day M, Muñiz LC, Bitran D, Arias R, Revilla-Sanchez R, et al. Activation of estrogen receptor-beta regulates hippocampal synaptic plasticity and improves memory. Nat Neurosci. 2008;11:334–343.

70. Bramham CR, Worley PF, Moore MJ, Guzowski JF. The Immediate Early Gene Arc/Arg3.1: Regulation, Mechanisms, and Function. Journal of Neuroscience. 2008;28:11760–11767.

71. Nikolaienko O, Patil S, Eriksen MS, Bramham CR. Arc protein: a flexible hub for synaptic plasticity and cognition. Seminars in Cell & Developmental Biology. 2018;77:33–42.

72. Béïque J-C, Lin D-T, Kang M-G, Aizawa H, Takamiya K, Huganir RL. Synapse-specific regulation of AMPA receptor function by PSD-95. Proceedings of the National Academy of Sciences. 2006;103:19535–19540.

73. Bats C, Groc L, Choquet D. The Interaction between Stargazin and PSD-95 Regulates AMPA Receptor Surface Trafficking. Neuron. 2007;53:719–734.

74. Swendsen J, Le Moal M. Individual vulnerability to addiction. Ann N Y Acad Sci. 2011;1216:73–85.

75. Egervari G, Ciccocioppo R, Jentsch JD, Hurd YL. Shaping vulnerability to addiction - the contribution of behavior, neural circuits and molecular mechanisms. Neurosci Biobehav Rev. 2018;85:117–125.

76. Nalberczak-Skóra M, Pattij T, Beroun A, Kogias G, Mielenz D, de Vries T, et al. Personality driven alcohol and drug abuse: New mechanisms revealed. Neuroscience & Biobehavioral Reviews. 2020;116:64–73.

77. Vandegrift BJ, You C, Satta R, Brodie MS, Lasek AW. Estradiol increases the sensitivity of ventral tegmental area dopamine neurons to dopamine and ethanol. PLOS ONE. 2017;12:e0187698.

78. Torres Irizarry VC, Feng B, Yang X, Patel N, Schaul S, Ibrahimi L, et al. Estrogen signaling in the dorsal raphe regulates binge-like drinking in mice. Transl Psychiatry. 2024;14:122.

79. Golden CEM, Martin AC, Kaur D, Mah A, Levy DH, Yamaguchi T, et al. Estrogen modulates reward prediction errors and reinforcement learning. Nat Neurosci. 2025;28:2502–2514.

80. Woolley CS. Acute Effects of Estrogen on Neuronal Physiology. Annu Rev Pharmacol Toxicol. 2007;47:657–680.

81. Smejkalova T, Woolley CS. Estradiol acutely potentiates hippocampal excitatory synaptic transmission through a presynaptic mechanism. J Neurosci. 2010;30:16137–16148.

82. Clements L, Harvey J. Activation of oestrogen receptor α induces a novel form of LTP at hippocampal temporoammonic-CA1 synapses. Br J Pharmacol. 2020;177:642–655.

83. Kim HJ, Casadesus G. Estrogen-mediated effects on cognition and synaptic plasticity: what do estrogen receptor knockout models tell us? Biochim Biophys Acta. 2010;1800:1090–1093.

84. Janak PH, Tye KM. From circuits to behaviour in the amygdala. Nature. 2015;517:284–292.

85. Grove-Strawser D, Boulware MI, Mermelstein PG. Membrane estrogen receptors activate the metabotropic glutamate receptors mGluR5 and mGluR3 to bidirectionally regulate CREB phosphorylation in female rat striatal neurons. Neuroscience. 2010;170:1045–1055.

86. Kumar A, Bean LA, Rani A, Jackson T, Foster TC. Contribution of estrogen receptor subtypes, ERα, ERβ, and GPER1 in rapid estradiol-mediated enhancement of hippocampal synaptic transmission in mice. Hippocampus. 2015;25:1556–1566.

87. Meitzen J, Mermelstein PG. Estrogen receptors stimulate brain region specific metabotropic glutamate receptors to rapidly initiate signal transduction pathways. J Chem Neuroanat. 2011;42:236–241.

88. Waters EM, Mitterling K, Spencer JL, Mazid S, McEwen BS, Milner TA. Estrogen receptor alpha and beta specific agonists regulate expression of synaptic proteins in rat hippocampus. Brain Res. 2009;1290:1–11.

89. Mühle C, Barry B, Weinland C, Kornhuber J, Lenz B. Estrogen receptor 1 gene variants and estradiol activities in alcohol dependence. Progress in Neuro-Psychopharmacology and Biological Psychiatry. 2019;92:301–307.

90. Cortes LR, Sturgeon H, Forger NG. Sexual differentiation of estrogen receptor alpha subpopulations in the ventromedial nucleus of the hypothalamus. Horm Behav. 2023;151:105348.

91. Hoffmann S, Gerhardt S, Mühle C, Reinhard I, Reichert D, Bach P, et al. Associations of Menstrual Cycle and Progesterone-to-Estradiol Ratio With Alcohol Consumption in Alcohol Use Disorder: A Sex-Separated Multicenter Longitudinal Study. Am J Psychiatry. 2024;181:445–456.

